# Self-organization of Drosophila chromatin architecture in a cell-free system

**DOI:** 10.64898/2026.03.17.712299

**Authors:** Muhunden Jayakrishnan, Gizem Kars, Aline Campos-Sparr, Magdalena A. Karpinska, Sanyami Zunjarrao, Noura Maziak, Carla E. Margulies, Juan M. Vaquerizas, Maria Cristina Gambetta, A. Marieke Oudelaar, Peter B. Becker

## Abstract

Metazoan genomes are organized by folding of the nucleosome fiber into loops and domains that support long-range regulatory interactions. Although cohesin-mediated loop extrusion and architectural DNA-binding proteins are central to current models of genome organization, how these mechanisms integrate to generate higher-order structure remains incompletely understood.

Early *Drosophila melanogaster* embryogenesis provides a unique window into the emergence of chromatin architecture, as rapid syncytial nuclear divisions occur largely in the absence of transcription. However, probing the mechanisms underlying this primordial folding in vivo is technically challenging.

Here, we establish an in vitro system that reconstitutes complex chromatin using extracts from syncytial embryos. Nucleosome mapping and Micro-C analyses reveal that long-range interactions, including loops and topologically associating domains (TADs), emerge spontaneously from soluble extract components. While some structures resemble those observed in early embryos, others represent latent interaction potentials that are constrained in vivo.

Focusing on the *eve* locus, we find that TAD formation is incompatible with a simple loop extrusion model and instead requires direct pairing of boundary elements mediated by the insulator protein ‘Suppressor-of-hairy-wing’ Su(Hw). Together, our work demonstrates that key features of 3D genome organization can be reconstituted in a cell-free system and provides a tractable platform for mechanistic dissection of chromatin folding in *Drosophila*.

## Main

Eukaryotic chromosomes consist of single, long DNA molecules that are organized as chains of nucleosomes, the repeating unit of chromatin. The wrapping of DNA around histone octamers to form nucleosome arrays can be visualized as ‘beads-on-a-string’ by electron microscopy in low-salt conditions. In interphase nuclei, however, these nucleosome arrays are hierarchically folded across multiple length scales, giving rise to chromosome territories, compartments, and fine-scale interaction patterns that are closely linked to genome regulation^1,2^.

A central feature of chromatin architecture is its non-random spatial organization. Polymer dynamics leads to stochastic encounters between chromatin segments, while short-range contacts within the fiber may be stabilized by electrostatic interactions and nucleosome packing, resulting in fiber conformations of varying geometry^3–7^. Longer-range contacts can be stabilized by dedicated architectural proteins, either through oligomerization or multivalent interactions^8–11^. Alternatively, two DNA segments may be transiently captured by a single protein complex, as proposed for the ‘loop capture’ mechanism of cohesin^12^.

From the structural perspective, such long-range contacts define loops of the chromatin fiber that are topologically isolated from neighboring chromatin^13^. Loop formation and long-range contacts may be actively promoted through loop extrusion by members of the SMC complex family, such as cohesin and condensin. In this mechanism, SMC complexes bind to DNA and reel the flanking nucleosome fiber into a loop via an ATP-dependent spooling reaction^14^ . Loop formation is halted at stably bound ‘barrier’ proteins like CTCF, which stall extrusion in an orientation-dependent manner^15–17^. Loop extrusion compacts the chromatin fiber, and in general segments within a loop are more likely to interact with each other than with sequences outside the loop. This is visualized in chromosome conformation capture assays as Topologically Associated Domains (TADs), the boundaries of which in mammals usually coincide with CTCF binding sites. Loop formation facilitates specific long-distance interactions of regulatory elements within the same TAD, giving rise to nested loops and sub-TADs^18^. These architectural principles facilitate and constrain functionally important long-range interactions, such as contacts between enhancers and promoters^19–21^, or between immunoglobulin heavy chain recombination sites^22,23^.

However, these principles are not universally conserved across metazoans. In *Drosophila*, CTCF contributes to only a subset of boundaries, while many TAD boundaries and loop anchors are occupied by other architectural proteins, including ‘suppressor-of-hairy-wing’ [Su(Hw)], BEAF-32, Pita, ZW5, CP190, GAF, and CLAMP^8,24–27^. Boundaries in flies frequently coincide with active promoters and regions of divergent transcription^28–30^, suggesting that multiple, context-dependent mechanisms such as transcriptional activity, nucleosome organization, and architectural protein clustering contribute to genome folding^26^. Whether these factors act primarily by stalling cohesin, by directly pairing boundary elements, or through alternative mechanisms remains an open question^16,31^.

The principles underlying the intricate interphase chromosome organization likely arise from multiple layers of organization^13^. The above principles suggest that a first level of organization of a naïve chromatin fiber could emerge from polymer dynamics and nucleosome interactions. A second level is imposed by functional interactions of loop extrusion complexes, barrier proteins and protein complexes bound at regulatory elements, such as promoters and enhancers^19,21,32,33^. A third level of organization arises from the confinement of the chromatin fiber in the limited space of a cell’s nucleus. Nuclear lamina and pores, nucleoli and other membrane-less compartments contribute surfaces to which certain types of chromatin are tethered^34^.

The early phases of *Drosophila* embryogenesis provide an exceptional experimental system to study the gradual emergence of chromosome organization. After fertilization, the embryo develops as a syncytium with 13 rapid divisions of nuclei. Because the G-phases of the cell cycle are lacking, the assembly of nuclei almost entirely relies on maternal stores of chromatin proteins^35,36^. Although a minor wave of zygotic transcription can already be detected as early as nuclear cycle (NC) 10, the majority of transcription is activated at NC14 in the cellular blastoderm stage, when nuclei have migrated to the embryo surface and cellularization happens. This is also the time when the linker histone H1 is incorporated into chromatin, constitutive heterochromatin forms and polycomb proteins establish facultative heterochromatin^37^. These early nuclei lacking H1 are 10-times larger than nuclei of later stages revealing the immature levels of genome folding^38^.

Monitoring the maturation of genome architecture during early embryonic development allows investigating the folding of the genome in the absence of widespread transcription. The genome architecture is mature at NC14, when transcription is high^28,39^. More recent Micro-C studies with enhanced sensitivity and resolution revealed that first TAD boundaries and loops form prior to zygotic gene activation (before NC9) and reveal the instructive role of insulator proteins and pioneer transcription factors, such as CTCF/CP190, Su(Hw), Pita, BEAF-32, ZW-5, GAF and Zelda^25,26^. These findings suggest that aspects of 3D genome organization can precede, and potentially scaffold, transcriptional activation.

Mistelli described chromatin as a self-organizing system^2^. Indeed, using recombinant chromatin and purified remodelers, Oberbeckmann et al., demonstrated that regularly spaced nucleosome arrays surrounding transcription factor binding sites are sufficient to generate domain-like interaction patterns in reconstituted yeast chromatin, even in the absence of transcription or loop extrusion^40^. These findings raise the possibility that intrinsic chromatin properties contribute significantly to genome folding, motivating complementary bottom-up approaches.

Here, we extend this self-assembly framework by reconstituting chromatin folding in a cell-free system derived from syncytial blastoderm *Drosophila* embryos, allowing us to probe metazoan genome organization under controlled conditions. Chromatin assembly extracts (henceforth referred to as DREX) are derived from the cytoplasm of *Drosophila* embryos within the first 90 minutes after egg laying. These extracts contain all components required to assemble added DNA into complex chromatin^41^. These extracts are rich in ISWI-type nucleosome sliding activities, such as ACF, NURF and CHRAC, that endow the reconstituted chromatin with dynamic plasticity^42^. The extract has been used to reconstitute chromatin on plasmids, fosmids and genomic DNA^43^. DREX-assembled chromatin mimics the naive ground state of the pre-ZGA genome: it lacks histone H1 and later-stage transcription factors or abundant histone modifications^44,45^ and is not transcriptionally active^41^. Long DNA fragments are assembled in nucleosome arrays with physiologically spacing that form condensates of physiological chromatin density^46^.

We previously discovered that DREX contains the Su(Hw) insulator complexes that bind to hundreds of sites in reconstituted genomic chromatin and serve as boundaries for nucleosome phasing^46,47^. Encouraged by this observation, we explored to which extent complex spatial genome organization patterns emerge in this cell-free system. We reconstituted syncytial blastoderm chromatin on a pool of BACs/fosmids representing ∼0.8% of the *Drosophila* genome and probed for long-range folding using nucleosome-resolution Micro-C and MNase-seq. This revealed complex folding patterns of the chromatin fiber, such as loops and TADs of sizes up to 35 kb, with the most prominent structure being the ‘volcano’ TAD at the *even-skipped (eve)* locus. Many chromatin loops are anchored at boundaries defined by binding of the Su(Hw) insulator. Some, but not all, structures resemble the corresponding long-range interactions that emerge during the nuclear cycles (NC) 9-14 of embryonic development.

The cell-free chromatin reconstitution system lends itself to mechanistic dissection of the reconstituted processes and structures. Focusing on the *eve* TAD, we manipulated the arrangement of boundary elements, levels of diagnostic protein factors and energy availability to explore the mechanistic basis of TAD formation. Our data are inconsistent with a cohesin-based loop extrusion mechanism and suggest the direct pairing of the Homie and Nhomie boundary elements mediated minimally by Su(Hw).

The assembly of complex chromatin in extracts of syncytial blastoderm embryos provides a powerful experimental approach for the exploration of long-range chromatin interactions in *Drosophila*, complementing more established ‘in vivo’ procedures. Our study provides a proof-of-principle that long-range chromatin folding can be reconstituted in vitro from soluble components.

## Results

### An in vitro system to study folding principles of complex reconstituted chromatin

To study 3D genome organization patterns of in vitro-reconstituted chromatin, we adapted a previously established system to reconstitute *Drosophila melanogaster* (‘*Drosophila’* henceforth) chromatin using cytoplasmic extracts derived from 0-90 minute-old syncytial blastoderm fly embryos (DREX) (Figure 1a). DREX-assembled chromatin mimics the genomic organization of embryos before the functional stratification that accompanies the Zygotic Genome Activation (ZGA). It lacks transcription factors and the transcription machinery as well as abundant histone modifications^41,44^, however it has a complex chromatin-associated proteome of ∼900 proteins^45^ including chromatin remodelers which were recently implicated in driving 3D genome organization patterns in yeast chromatin in vitro^40^. As a consequence, reconstituted chromatin contains well-positioned nucleosomes and nucleosome free regions (NFRs) predominantly at non-TSS sites (consistent with lack of transcription) marked by sequence motifs for the Suppressor of Hairy Wing [Su(Hw)] insulator complex and Phaser silencer complex^47^. Further, DREX-assembled chromatin also spontaneously forms micron-scale condensates, indicative of emergent higher order chromatin interactions^46^. These prior observations motivated us to investigate the folding patterns of DREX-assembled chromatin using high-resolution Micro-C.

**Figure 1.**
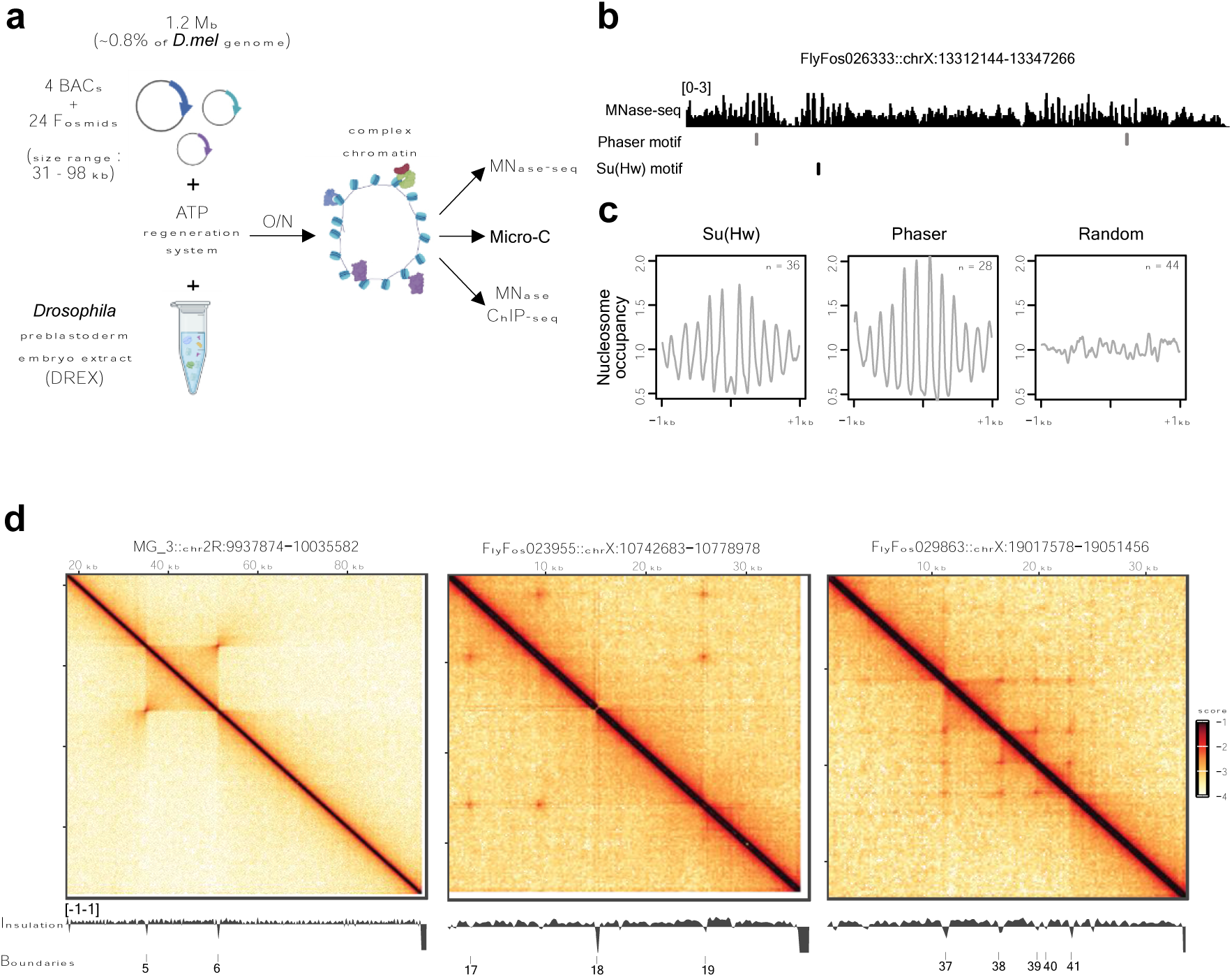
**Assembly of complex, folded chromatin in extracts of syncytial blastoderm *Drosophila* embryos.** a) Schematic illustrating the experimental protocol and key features of the in vitro chromatin assembly reaction using DREX and subsequent assays. Illustration created with Biorender.com. O/N: overnight. b) MNase-seq data indicating nucleosome positions in DREX-assembled chromatin. The fosmid identity as well as original dm6 coordinates are indicated above. The positions of Phaser and Su(Hw) motifs are marked below the MNase-seq track. c) Composite plots representing average MNase-seq signal in 2-kb windows centered at Su(Hw) and Phaser motifs. The number of motifs is indicated by n. Randomly centered coordinates serve as a negative control. Coverage values for individual sites are provided in source data. d) Micro-C contact maps for representative BACs at 200 bp resolution. The dm6 chromosomal coordinates of the cloned *Drosophila* sequences are indicated above the panels. The distance from insert start is indicated above the heatmap. The scale bar to the right applies to all panels. Diamond insulation scores as well as called boundaries with index numbers (see text) are indicated below the heatmap for reference.

To generate very high coverage contact maps, we resorted to a reduced representation of the *Drosophila* genome by an equimolar pool of non-contiguous cloned DNAs: 24 fosmids representing X chromosomal sequences (median insert size 35 kbp) and 4 BACs containing DNA derived from chromosome 2R (median insert size 90 kbp), totalling ∼1.2 Mbp (∼0.8% of the *Drosophila* genome). These constructs were selected based on various features detailed in ‘Methods’. Incubation of this DNA pool with DREX and an ATP regeneration system results in chromatin assembly with physiologically spaced nucleosomes, assessed by MNase-seq coverage around representative Su(Hw) and Phaser motifs (Figure 1b). Cumulative plots representing average MNase-seq signal at Su(Hw) and Phaser sites confirm the generation of NFRs at motif centers as well as phased nucleosome arrays (PNAs) around the motifs, while no such patterns were found at 44 random coordinates (Figure 1c).

We then applied a modified version of the in vitro Micro-C protocol to investigate the folding patterns of DREX-associated chromatin. To minimize variability from different DREX batches, we pooled multiple independent experiments to generate a dataset with ∼70 million valid contacts for the 1.2 Mbp *Drosophila* sequence. We visualized the resulting contact maps at 200 bp resolution, which revealed several examples of complex folding patterns including TADs and loops (Figure 1d). To facilitate domain identification, contact maps were complemented with insulation profiles, which were used to call 44 boundaries (which frequently coincided with loop anchors).

An example of a complex structure includes a 16-kb ‘volcano’ TAD, reconstituted at the even-skipped (*eve*) locus (BAC ID MG_3), which had been previously described to occur based on a specific-pairing interaction of the Nhomie-Homie insulator elements flanking the *eve* gene^11,48,49^. Contact maps of all 28 constructs alongside reference insulation profiles, Phaser and Su(Hw) motifs, and ChIP-seq peaks for Zelda (a pioneer transcription factor) from 2-4 hr embryos are shown in Figure S1. Of note, we do not observe any prominent trans-interactions occurring across constructs. (Figure S2).

Other examples of complex structures include ‘mini’ TADs spanning only a few kb (1.8 kb TAD in BAC ID MG_1), occurrences of isolated insulation dips in the absence of paired boundary interactions required to form a canonical TAD (FlyFos022038) and instances of nested loops (FlyFos029863). Overall, ∼50% of the constructs displayed some level of 3D genome organization. Taken together, these results show that complex nucleosomal and genome-folding patterns can be reconstituted from soluble components outside the complex milieu of a nucleus.

#### Protein factor occupancy correlates with 3D genome organization patterns in DREX chromatin

A previous study, which examined chromatin folding patterns in salt-gradient dialyzed chromatin remodelled by yeast TFs and remodelers, demonstrated that NFRs and PNAs were sufficient to reconstitute chromatin domains^40^. Given that DREX contains several nucleosome positioning and remodelling factors, we sought to verify if similar principles applied to our system (Figure S3). To identify PNAs in an unbiased fashion, we used our previously published approach^47^ to scan for regions of high periodicities in MNase-seq data. Applying this approach, we identified 122 PNAs, which we further stratified into Su(Hw)-associated, Phaser-associated or other non-insulator types (characterized by low complexity sequences with intrinsic capability to position nucleosomes). To understand the correlation between nucleosome positioning behaviour and Micro-C insulation, we visualized a 17 kb representative region on a fosmid containing the three classes of PNAs in close proximity (Figure S3a). Only the PNA associated with the Su(Hw) motif correlated with a prominent dip in insulation score (indicated by a red asterisk), in addition to participating in a looping interaction with downstream Su(Hw) motifs. To quantify this system-wide, we calculated the average spectral density estimate (SDE), which captures nucleosome array regularity, together with Micro-C insulation scores at these regions and visualized them as boxplots, which revealed that only Su(Hw)-associated PNAs showed more negative insulation scores (i.e. strong boundaries), while Phaser and non-insulator sites did not function as boundaries, despite high array regularity (Figure S3b). These observations argue against a straightforward relationship between nucleosome positioning and 3D genome organization in this system.

We then sought to understand the potential protein factor composition at the 44 boundaries by integrating published ChIP-seq datasets from *Drosophila* cell lines and developing embryos, recognizing that these data reflect in vivo binding and do not necessarily imply factor presence or occupancy in DREX-assembled chromatin. The resulting clustered heatmap, which represents average factor occupancy at each boundary, revealed interesting subclasses of boundaries (Figure 2). We complemented the clustered heatmap by additionally visualizing direct Su(Hw) binding in DREX by MNase ChIP-seq as well as ATAC-seq accessibility from Nuclear Cycle (NC) 11-13 embryos. Strikingly, around 50% of the boundaries appeared to be correlated to strong Su(Hw) and moderate Cp190 in vivo occupancies alone. These include example boundaries #38, 39 and 41 located on FlyFos029863, which also form a network of looping interactions (Figure 1d, Figure S1). Interestingly, the upstream boundary #37 shows an additional potential enrichment of Zelda and Pita. The other major subclasses of in vitro-detected boundaries show strong enrichment for many ‘active’ factors, such as promoter-binding factor BEAF-32 and RNA Pol II. These sites generally show a lower enrichment of Su(Hw) binding in vivo, and are largely unbound in vitro, suggesting that Su(Hw) is possibly recruited indirectly (i.e., through interactions with other factors) to these sites. Further, these sites also frequently display high accessibility in vivo, which may be related to regulatory elements like promoters and enhancers. Examples of these types of boundaries include #17-19 located on FlyFos023955, which also function as loop anchors (Figure 1d, Figure S1). The last major subclass does not show an enrichment for any factor (bottom third of the heatmap), and likely includes a combination of weak boundaries (for instance, #28 located within FlyFos019248) and false-positive boundary calls (for instance, #36 located within FlyFos025442). Importantly, while these subclasses describe broad patterns, multiple cases of unique, locus-specific factor compositions are observed. Most notably, boundaries #5 and #6 flanking the *eve* locus show moderate-to-strong enrichment of most factors interrogated in this study, including Su(Hw) and Rad21 (component of the cohesin complex). Collectively, these observations argue for emergent, protein-directed genome organization in DREX-assembled chromatin.

**Figure 2.**
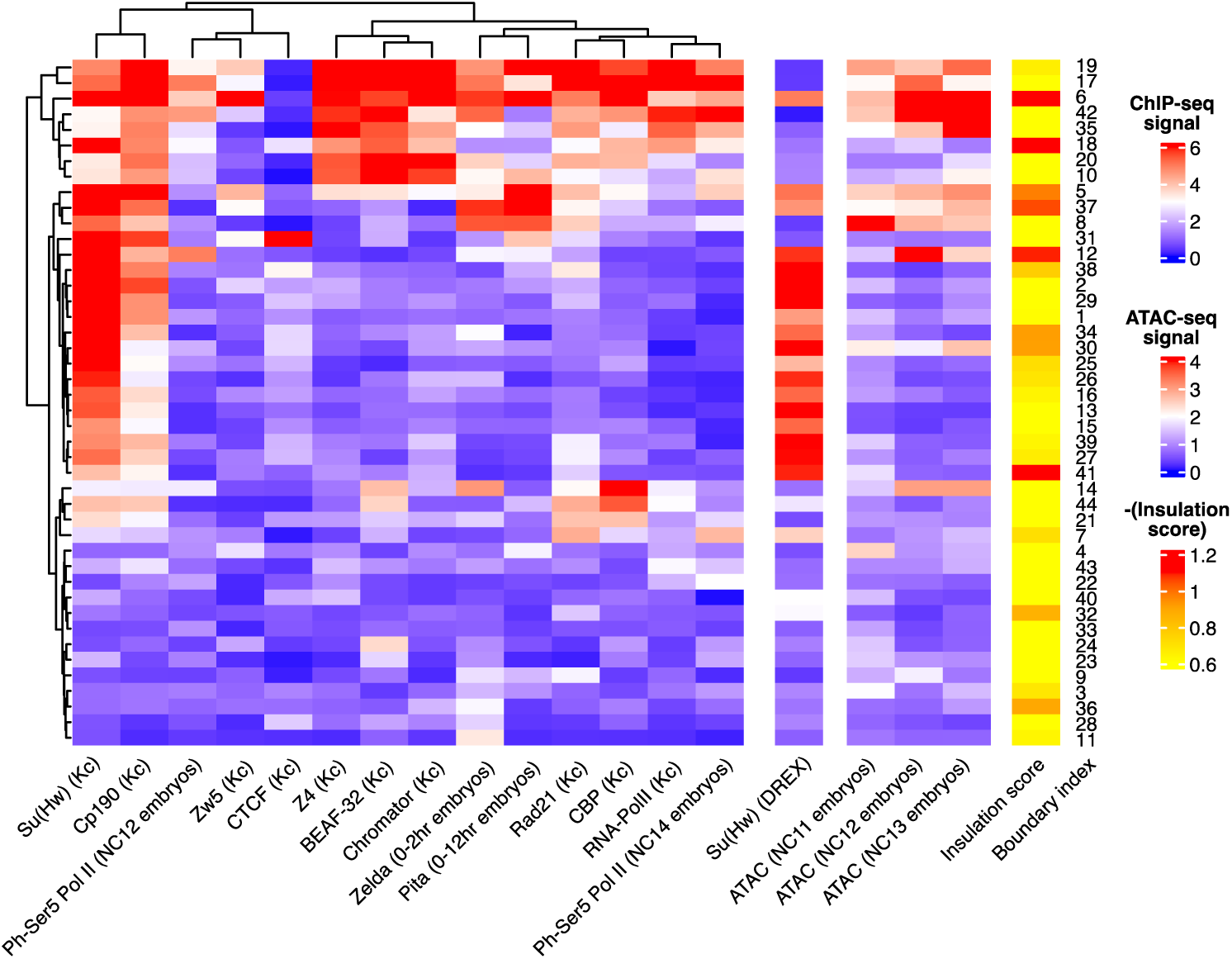
Putative chromatin-binding factor composition at insulating in vitro boundaries. Hierarchically clustered heatmap of average factor ChIP-seq enrichment in 1-kb windows around 44 in vitro boundaries. Each row represents a boundary (indicated by numerical index on the right side of the map) and each column represents ChIP-seq enrichment score for a given factor. ChIP-seq target as well as model system is indicated below each column. Kc refers to the Kc167 cell line, and developmental ages in hours of embryo-derived ChIP-seq datasets are indicated. Direct Su(Hw) occupancy in DREX-reconstituted chromatin, ATAC-seq for NC11-13 embryos, and negated in vitro insulation scores (plotted such that higher values correspond to stronger boundary insulation) for the 44 in vitro boundaries ordered according to the main clustered heatmap are provided for reference. Heatmap color scale bars are shown on the right.

#### In vitro structures often do not resemble in vivo folding patterns

While early HiC experiments had suggested that genome folding patterns are largely established concomitant with major ZGA (NC14) in *Drosophila*^39^, more recent studies utilizing higher-resolution Micro-C have shown that elements of higher-order genome organization, including many TAD boundaries, are already detectable at earlier stages of embryogenesis^25,26^. To assess whether and when interactions observed in vitro also occur in vivo, we drew on recently published Pico-C contact maps along with in vivo boundary classes of stage-resolved *Drosophila* embryos (Figure 3). We anticipated that the in vitro structures would closely mirror the in vivo folding landscape of pre-ZGA embryos, however, our results unexpectedly uncovered striking discrepancies between the two contexts. We broadly grouped the observed structures as ‘in vivo only’, shared, or ‘in vitro only’ based on the (dis-)similarity of their contact profiles in vitro and in vivo (Figure 3a).

**Figure 3.**
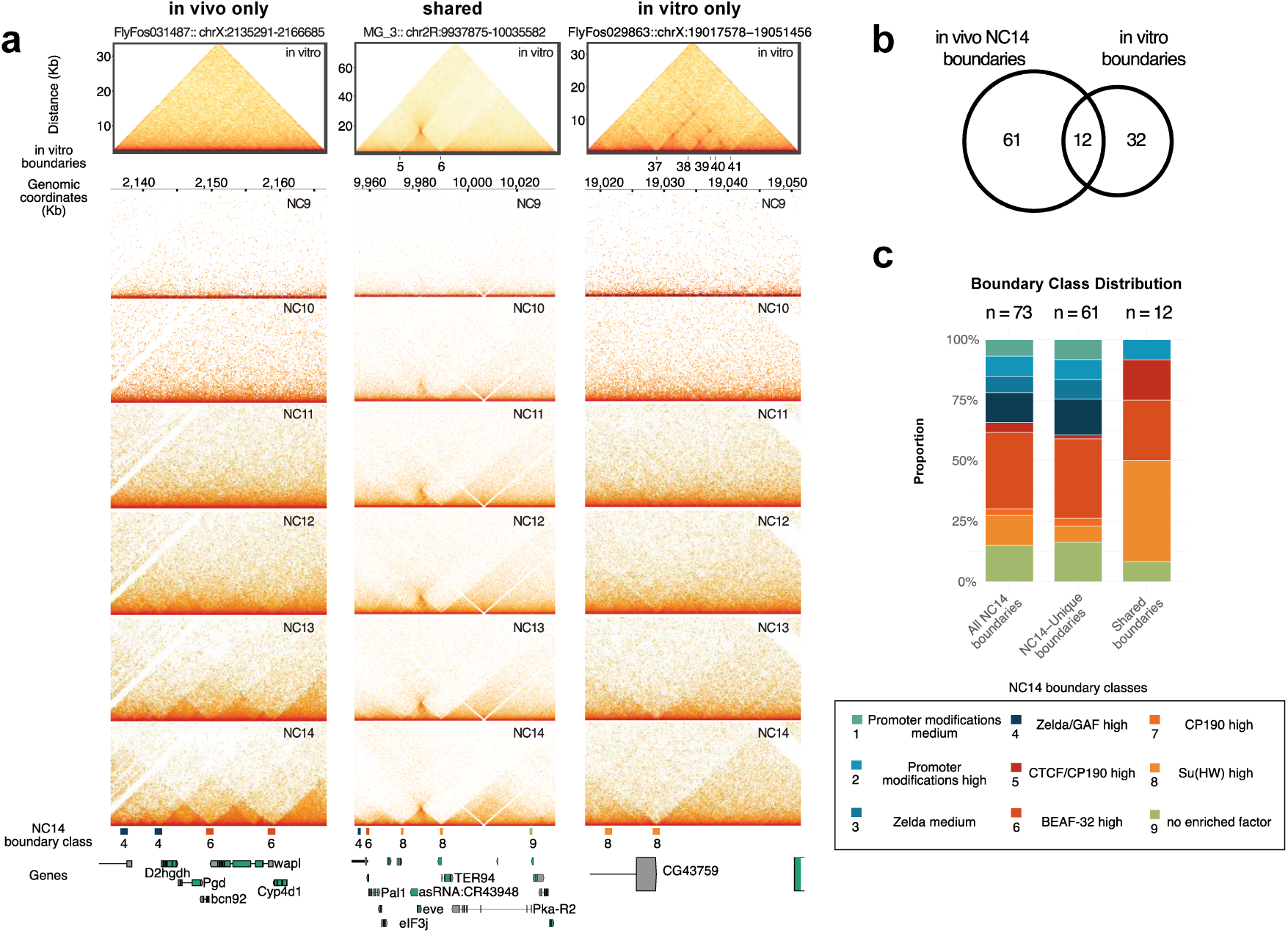
**Comparison between 3D chromatin folding patterns in vivo and in vitro** a) Top: 200-bp resolution horizontal Micro-C contact maps for representative DREX-chromatinized regions for three groups (‘in vivo only’, ‘shared’, ‘in vitro only’) defined based on correlation with in vivo data. Region identifiers are indicated above. Y-axis values indicate distance from insert start. Identified in vitro boundaries along with reference indices are shown below the contact heatmap. Below: 125-bp resolution Pico-C contact maps from developing *Drosophila* embryos for in vitro-matched genomic regions. Developmental stages indicated by NC9-14 are shown in the top-right corners of the contact maps. Chromosomal coordinates (dm6) are indicated above. Representative genes, positions of boundaries identified in NC14 embryos and boundary classes are shown below NC14 contact maps. Brief descriptions of in vivo boundary classes are shown in the legend of (c). b) Venn diagram representing counts of distinct and shared boundaries (defined based on proximity within 350 bp) between in vivo NC14 and in vitro boundaries contained within the 1.2 Mb region focussed on in this study. c) Stacked barplots representing proportions of in vivo NC14 boundary classes 1-9 for all NC14 boundaries (left), NC14-only boundaries (middle) and boundaries shared between NC14 embryos and in vitro DREX chromatin (right). The number of boundaries for each group is indicated above the stacked bars. Numerical counts for each class are provided in source data. Brief descriptions of the in vivo boundary classes are shown in the legend below the main plot.

Some regions, such as those contained within FlyFos031487, are largely unstructured in vitro but demonstrate clear domain organization in vivo (Figure 3a, left column). Frequently, these regions acquire discernible structures in vivo only close to the onset of ZGA. Examination of the in vivo boundaries within FlyFos031487 (see track below contact maps) revealed they are of Types 4 (Zelda and GAF high) and 6 (BEAF-32 high active promoters). The simplest explanation for the absence of these structures in vitro is that DREX lacks the instructive factors, such as GAF^46^ and active transcription^41^.

Other in vitro structures, for instance the *eve* TAD, resembled the in vivo interaction profiles (Figure 3a, middle column). The Nhomie-Homie loop is detectable as early as NC9 embryos, with the TAD progressively getting stronger over subsequent cycles. These in vivo boundaries are classified as Su(Hw)-enriched Type 8 boundaries, consistent with the observed enrichment of Su(Hw) at boundaries #5,6 in vitro.

Interestingly, we observe several instances where loops are formed in vitro but do not have an in vivo correlate. For example, loop interactions anchored by Su(Hw) on FlyFos029863 do not occur at any developmental stage in vivo (Figure 3a, right column). This region forms loops in vitro and the Su(Hw)-clustered boundary participates in multiple long-range loops in vivo, but not at corresponding positions. Interestingly, one of the in vitro loop anchors (#37) instead acts exclusively as a strongly insulating Type 8 Su(Hw)-occupied boundary in NC14 embryos. Further, in vitro boundaries #38,39 and 41 show strong Su(Hw) binding in both model systems, but do not correspond to boundaries in nuclei, suggesting that the looping potential of Su(Hw) is constrained in vivo. We highlight several additional examples of in vitro-specific loops (FlyFos023955), in vivo-specific loops (FlyFos022038) and shared (FlyFos029355) in Figure S4.

To get a better understanding of the correlations system-wide, we compared the called in vitro boundaries with the boundaries obtained from NC14 embryos. Restricting our analyses to the regions contained within the BAC/Fos system resulted in 73 NC14 boundaries (82 prior to filtering for problematic blacklisted in vitro Micro-C regions), compared to only 44 in vitro boundaries (Figure 3b). The overlap analysis confirmed our initial qualitative observations; only 12 boundaries were shared between the two contexts. A comparison of boundary types represented in this group suggests that DREX is capable of reconstituting a subset of boundaries that are enriched in Su(Hw), BEAF-32 and CTCF/CP190 (Figure 3c). Evidently, not all boundaries of a single class are reconstituted in vitro, suggesting that subclasses with different requirements exist (for example, despite established Su(Hw) function in DREX, 4 out of 9 class 8 boundaries are not reconstituted).

In summary, these results show that while complex boundary types can be reconstituted in vitro, the cell-free system also reveals <u>potential</u> long-range chromatin interactions mediated by Su(Hw) that are not realized in vivo. We speculate that the selection of contacts from all possible combinations is influenced by either competitive constraints or requires levels of organization, such as chromosome anchoring at the lamina, that are missing in the extract.

#### Leveraging in vitro reconstitution to test loop extrusion versus boundary pairing

While cohesin-mediated loop extrusion has been established as a major mechanism by which the mammalian genome is packaged and regulated, the extent of loop extrusion as a principal mechanism of 3D genome organization in *Drosophila* is still unclear^16,31,50^. While recent studies supported a model involving direct pairing of boundary elements without the requirement for extrusion, using reporter assays and sequence manipulations^11,51^, direct evidence remains limited, in part because key extrusion-related factors are essential for development and therefore difficult to deplete. DREX-reconstitution offers an experimental approach to address this question. We focus on the *eve* locus for two reasons: first, its boundaries, Nhomie and Homie, are known to be bound by a complex compendium of factors, including Su(Hw) and Rad21, allowing to test the relative contributions of the direct pairing versus loop extrusion mechanisms. Further, the *eve* TAD is by far the strongest reconstituted structure and appears robustly even in shallowly-sequenced, low-complexity libraries, which simplifies experimental assessment. To distinguish between the two mechanisms, we designed three perturbations, each of which probes a different mechanistic aspect: 1) ATP depletion to explore the energy dependence of TAD maintenance; 2) diagnostic cleavage of the input DNA to change distance and orientation of the two boundary elements or to physically separate them prior to TAD formation; and 3) immunodepletion of critical protein factors involved in either mechanism prior to TAD establishment.

#### Active ATP hydrolysis in dispensable for maintenance of eve structure

Given that many chromatin regulators are ATP-dependent motors or remodelers and loop extrusion relies on continued ATP hydrolysis^52,53^, we asked whether sustained ATP availability is required to preserve established 3D genome architecture (Figure 4a). Because the assembly of regularly spaced, physiological chromatin in the DREX system requires ATP hydrolysis, we can only assess the energy dependence once chromatin assembly is complete. We first measured the ATP consumption kinetics during the course of chromatin assembly using CellTiter-Glo, a luminescence-based assay (Figure 4b) by diluting a small volume of the assembly reaction 500-fold to ensure that the measured luminescence values were within the linear sensitivity range of the assay. Measurement of plain DREX provided a base line.

**Figure 4.**
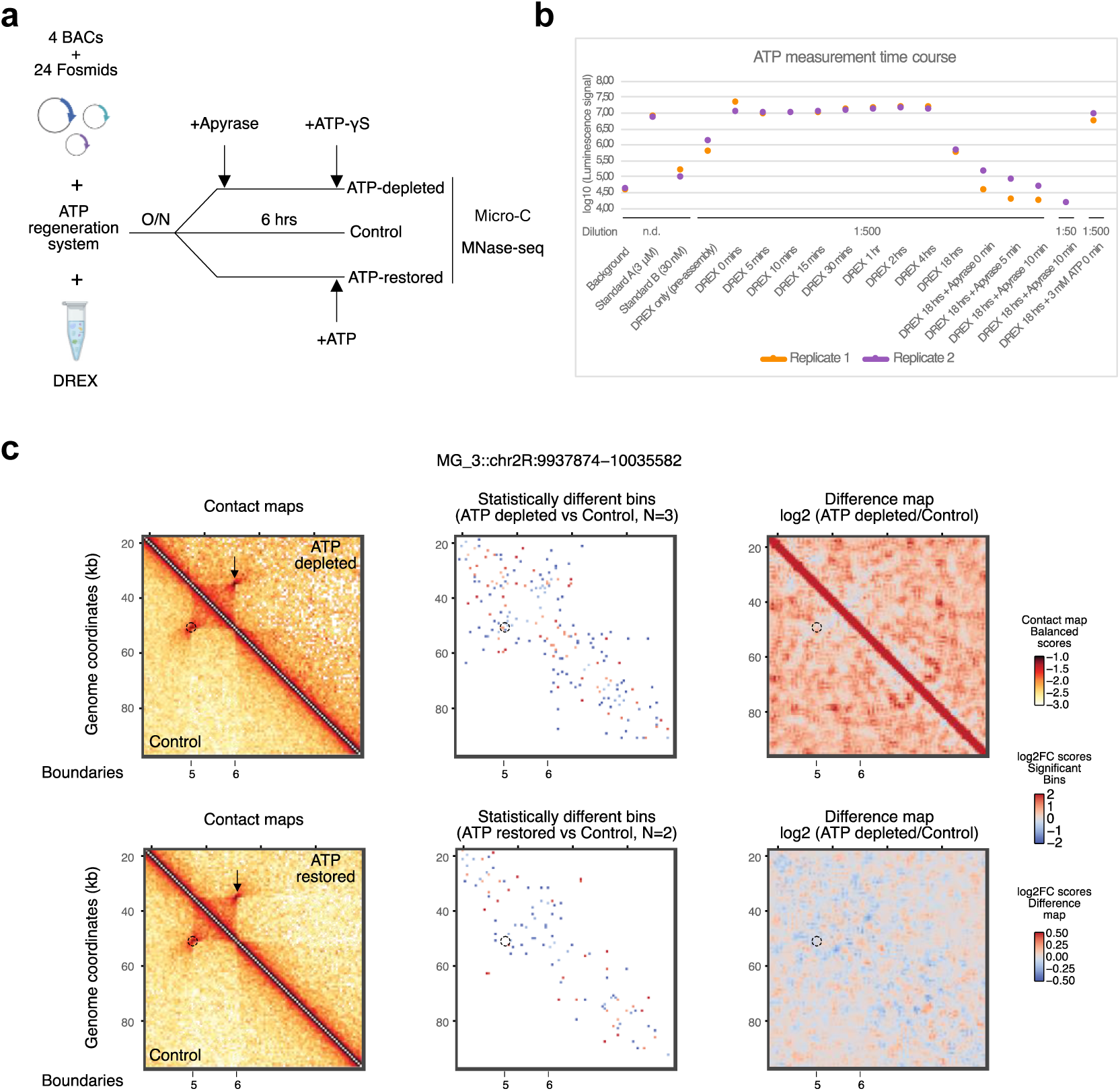
**Active ATP hydrolysis in dispensable for maintenance of the *eve* TAD** a) Schematic illustrating experimental timeline for manipulation of ATP levels in DREX assembly reactions and subsequent assays. Illustration created with Biorender.com b) Dot plot representing log10-transformed CellTiter-Glo luminescence measurements at different time points during the assembly reaction and following manipulations of ATP levels. The color of the points represents the associated biological replicate as indicated at the bottom. The dilution factor used for measurements (n.d. implies no dilution) as well as the timepoint/treatment labels are indicated below the scatter plot. c) 800-bp resolution Micro-C maps of the *eve* locus characterizing changes in contact patterns upon either ATP depletion (top row) or after ATP restoration (bottom row). For each treatment condition; Left: Interaction maps representing balanced contact frequencies for control (below the diagonal) or treated (above the diagonal) samples. Center: Differential bins (|log-fold change| >= 0.35 and p.value < 0.1) derived from statistical testing of indicated treatment versus control (with number of biological replicates used for testing denoted by N). Right: Difference maps representing log2-ratio of Treatment over Control contact frequencies. The dotted circle and arrow represent the Nhomie-Homie loop; called boundaries are denoted below. Scales for each heatmap type are shown on the right.

Supplementing the reaction with 3 mM ATP and an ATP regeneration system according to the DREX assembly protocol elevated the ATP levels 10-fold. Due to the regenerative system, these ATP levels remained high for the first 4 hours of chromatin assembly and maturation. At 18 hours, the ATP levels were reduced to the basal level of DREX, but still substantially higher than background (buffer only without ATP). ATP can be restored to initial levels by addition of 3 mM ATP, if needed (Figure 4b, last data point). To rapidly deplete residual ATP from overnight assemblies, we added the enzyme apyrase, which hydrolyzes nucleoside tri/diphosphates to AMP and inorganic phosphate. The measurement confirmed that by 10 minutes of apyrase treatment ATP levels drop to background levels. Measuring a more concentrated version of the depleted samples confirmed that residual ATP levels are well below the detection limit of the reagent. Together, these perturbations (ATP restoration and depletion) capture the entire range of ATP available during the assembly reaction.

We initially attempted to perform Micro-C with ATP-depleted samples, but apyrase treatment led to visible precipitation of chromatin, which resisted MNase digestion and ligated inefficiently (data not shown). We reasoned that ATP hydrolysis may lead to increased concentrations of Mg^2+^ ions, which are known to aggregate and precipitate chromatin in a concentration-dependent manner^54^. To avoid this unwanted side effect, we added back ATP-γS, a non-hydrolyzable variant of ATP, to partially restore the charge balance (Fig 4a,b). The resulting chromatin behaved more similarly to control chromatin in MNase digestions and ligations.

We first examined the MNase-seq profiles from the various perturbations, including short-term (10 min) and long-term (6 hour) ATP depletions with and without ATP-γS supplementation (Figure S5a). Visual inspection of the MNase-seq coverages, along with the periodicity (SDE) scores at various PNAs (indicated by blue boxes at the bottom), demonstrated that nucleosome positions were largely unchanged upon ATP manipulations. To quantify this response globally, we measured the mean SDE scores at the previously described PNA classes (including random regions as negative controls) in Control, 6 hours ATP-depleted (with ATP-γS supplemented), and ATP-restored chromatin (Figure S5b). The resulting violin plots revealed no significant differences for the different treatments for all PNA classes.

We then turned to the Micro-C data of the *eve* locus for the three conditions mentioned above (Figure 4c). We augmented the interaction frequency maps of treatment (left, above diagonal) and control (left, below diagonal) along with a map of statistically significant differential contacts (middle) and a log-ratio (treatment/control) difference map (right). Although the statistical difference maps contained sporadic significant pixels likely reflecting background noise, we primarily assessed for local clusters of contacts showing uniform directional changes. By this criterion, the *eve* TAD structure, including the Nhomie-Homie loop (Figure 4c, highlighted by dotted circle/arrow) was largely unchanged upon either ATP depletion or restoration of 3 mM ATP. Of note, we observed a general increase in background interaction frequencies upon ATP depletion, which may be explained by general compaction of chromatin in the presence of elevated Mg2+ concentrations, due to incomplete charge compensation. While our libraries generally weren’t complex enough to make robust conclusions about other structures, we observed other possibly interesting trends. For example, a 7-kb loop originating from boundary #17 on FlyFos023955 showed a slight weakening upon ATP depletion and a reciprocal strengthening upon ATP restoration. (Figure S5c).

We conclude that the *eve* TAD domain is maintained in the absence of ATP-dependent processes. We speculate that the depletion of ATP reduces chromatin dynamics, thus ‘freezing’ nucleosome positions and (some) long-range interactions.

#### Cleavage of DNA within the *eve* locus does not impair TAD formation

Sequence manipulation has been used to study the requirements for chromatin looping and TAD formation, but in vivo this requires cumbersome CRISPR-based approaches. The in vitro system lends itself to such experiments. The mechanism of loop extrusion requires that the fiber between the two loop anchors is continuous. Breakage of the fiber leads to disruption of the loop and should also compromise the TAD organization. To test the requirement for fiber continuity for *eve* TAD formation, we performed a series of restriction enzyme (RE) digestion experiments of the circular DNA pool prior to chromatin assembly (Figure 5a). The choice of the appropriate enzyme allows evaluating the importance of the intactness, distance and directionality of the boundary motifs for organizing the *eve* locus. A schematic of the locations of the different cut sites is shown in Figure 5b. The cleavage efficiency of DNA used in the assembly reactions was estimated by visualizing replicate MNase-seq profiles around the *eve* locus (Figure S6a). The positions of the cut-sites are indicated by asterisks (colors match the schematic in Figure 5b) and positions of boundaries #5 and 6 are provided for reference at the bottom. Visual inspection confirms the loss of sequencing coverage across the RE motifs upon digestion. Heatmaps representing MNase-seq signals 2-kb upstream and downstream of the cleavage sites confirm that most of the sequenced DNA molecules were cut (Figure S6b). In addition, sequences in the immediate vicinity of the cut site are lost (usually ∼1 kb), due to previously documented activity of end resection enzymes in DREX^55^.

**Figure 5.**
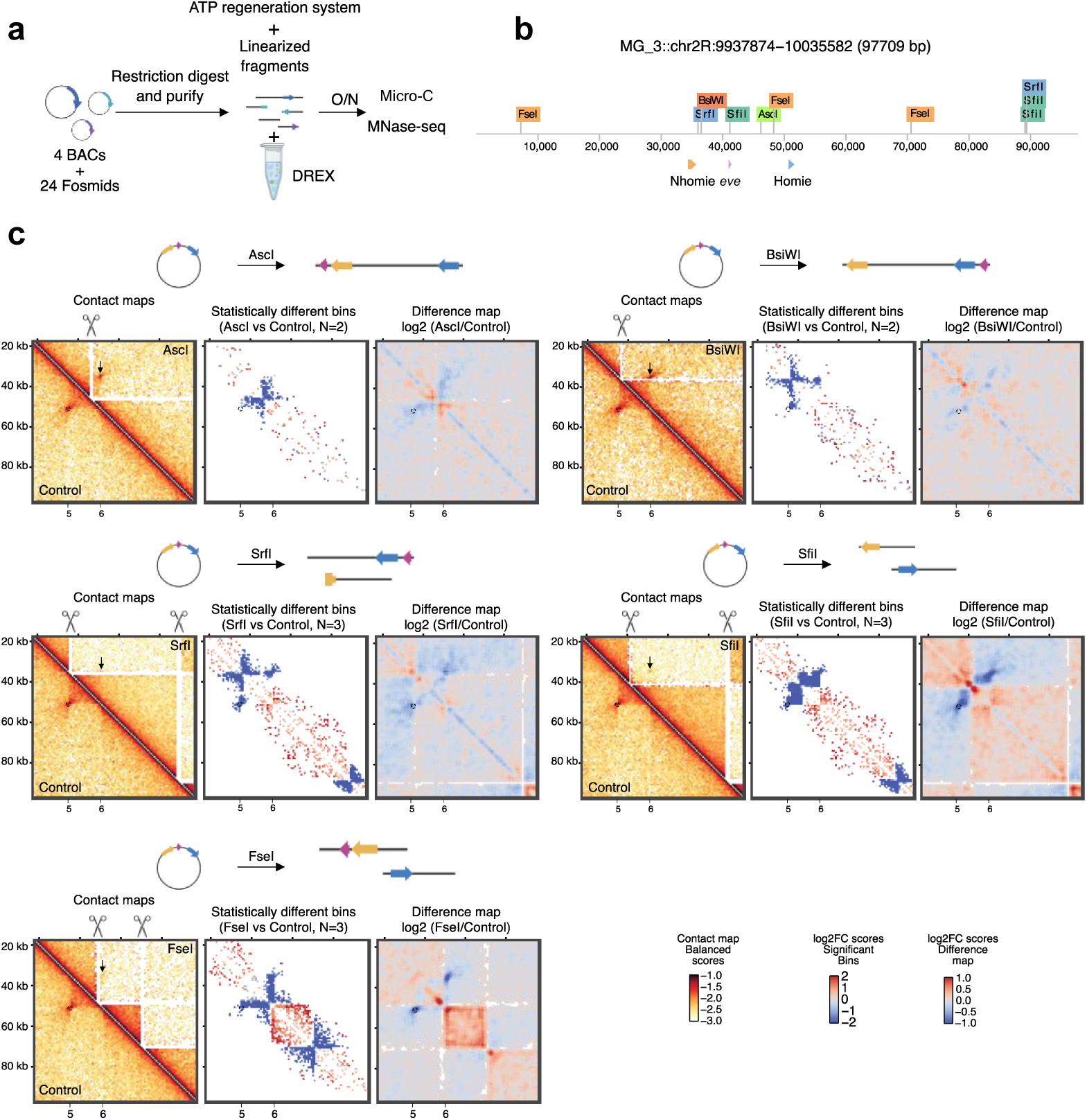
**Cleavage of constructs prior to assembly does not impair long-range interactions at the *eve* locus** a) Schematic illustrating experimental protocol for assaying the effect of DNA cleavage on chromatin interactions. Illustration created with Biorender.com b) Schematic of restriction digestion sites relative to the *eve* locus used in this study. c) 800-bp resolution Micro-C maps of the *eve* locus characterizing changes in contact patterns upon restriction digestion with single-cutters (AscI and BsiWI) or multi-cutters (SrfI, SfiI and FseI). For each restriction digestion condition; Left: Interaction maps representing balanced contact frequencies for control (below the diagonal) or digested (above the diagonal) samples. The expected cut site is denoted by a scissor icon. Center: Differential bins (|log-fold change| >= 0.35 and p.value < 0.1) derived from statistical testing of experimental condition versus control, as indicated (with number of biological replicates used for testing denoted by N). Right: Difference maps representing log2-ratio of digested over Control contact frequencies. The expected relative configuration of motifs on cut fragments are depicted above the maps. The dotted circle and arrow represent the Nhomie-Homie loop; called boundaries are denoted below. Shared scales for each heatmap type at the bottom of the figure panel.

Cleavage of the *eve* locus with either AscI (located about 4.5 kb upstream of Homie) or BsiWI (located about 800 bp downstream of Nhomie) (Figure 5c, top row) changes the relative arrangement of the boundary motif configurations, resulting in the *eve* gene being ‘outside’ the loop anchors, and increases the shortest distance between the loop anchors from ∼15 kb to ∼92 kb (assuming a 12.5 kb vector backbone). In addition to the normalized contact maps, we also visualized the differential interaction bins and log-ratio difference maps in a format similar to Figure 4c. In all cases, DNA cleavage leads to loss of ligated fragments originating around the cut-site. Cutting the *eve* locus with AscI leads to marginal reduction in internal interaction frequencies between Homie and the rest of the *eve* TAD, but the Nhomie-Homie loop as well as Nhomie-*eve* TAD internal interactions are formed remarkably well. Likewise, the loop and internal interactions are largely unaffected upon cleavage with BsiWI. In both experiments, although not identified as significant, a trend towards reduced contact frequencies of the ‘plume’ region (representing interactions between regions immediately upstream of Nhomie and downstream of Homie) is observed, which was previously proposed to depend on the relative orientation of the motifs^51^.

We then turned to the set of multi-cutters (Figure 5c, middle and bottom rows). The enzyme SrfI has one cut site ∼300 bp downstream of Nhomie and another site ∼39 kb downstream of Homie. The cuts lead to loss of the Nhomie motif due to resection, leaving a partially truncated sequence located in a fragment separate from the Homie motif and the *eve* gene. As expected, the loop interaction between Homie and Nhomie is no longer detected, but internal interactions between Homie and the rest of the *eve* TAD still occur. The other two enzymes (SfiI and FseI) leave both anchors intact, although they are now located on two separate DNA fragments, requiring ‘*trans*’ interactions. In both settings, the loop interaction is much weaker, although a very slight signal relative to the local background may still be observed. In addition, similar to all the previous cases, interactions between the *eve* proximal sequences with the remaining boundary element on the same DNA fragment can be observed.

We conclude that the Nhomie-Homie loop is remarkably unaffected by relocations *in cis* and may even interact very weakly *in trans*. Further, the Nhomie-Homie loop is separable from interactions of the individual boundaries with the *eve* proximal sequences, highlighted by SrfI ‘deletion’ experiment. These observations are at odds with a simple loop extrusion model involving the extrusion of DNA between the loop anchors and rather argue for a more complex interaction network involving Nhomie-Homie-*eve*.

#### Immunodepletion of cohesin and Su(Hw) argues against a loop extrusion model for the formation of the *eve* TAD

So far, all evidence against a cohesin-driven loop extrusion mechanism underlying the formation of the *eve* TAD, including the experiments presented above, has been indirect. If true, the *eve* TAD should form in the absence of cohesin. The depletion of cohesin in cells is difficult due to its vital functions in chromosome segregation, cell viability and embryonic development. In contrast, the cell-free system enables the immunodepletion of factors of interest from DREX prior to assembling chromatin to assess their importance for the establishment of chromatin fiber folding (Figure 6a). To test the contribution of loop extrusion by cohesin complexes versus direct boundary pairing by Su(Hw) in the establishment of *eve* structure, we immunodepleted the respective components (Rad21 and Su(Hw) respectively) from DREX with specific antibodies (Figure 6b). A non-specific IgG antibody served as a control to monitor the non-specific adsorption to the Dynabeads. Following magnetic separation of beads, the bead-bound proteins and the factor-depleted supernatant were analyzed by Western blotting. This confirmed an ∼80-90% reduction of the immunoprecipitation targets from DREX (Figure 6b, Figure S7a).

**Figure 6.**
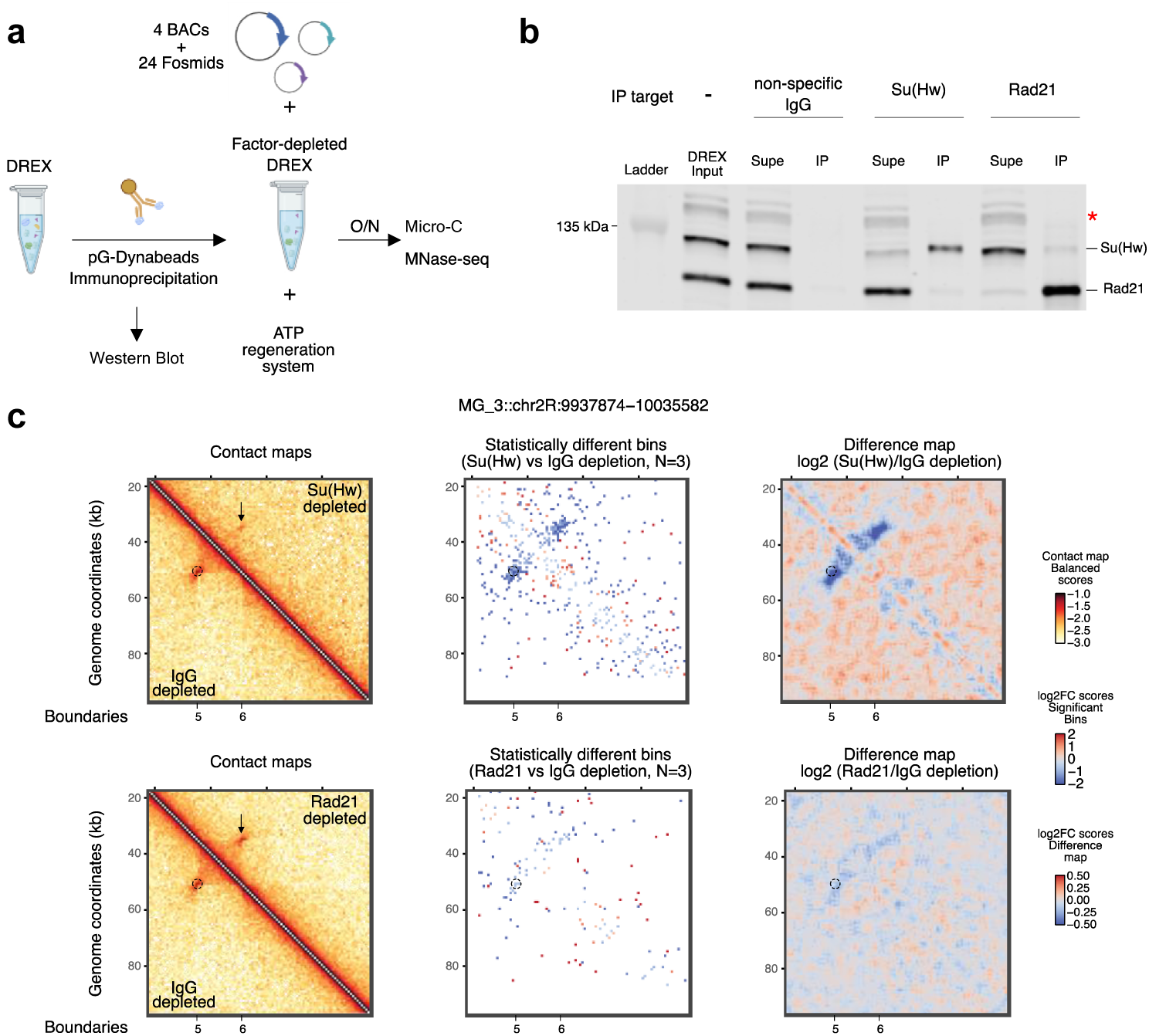
**Effects of immunodepletion on loop formation at the *eve* locus** a) Schematic illustrating experimental protocol for antibody-mediated factor depletion for DREX assembly reactions and subsequent assays. Illustration created with Biorender.com b) Representative Western blot for estimation of depletion efficiencies. ‘Input’ samples refer to plain DREX collected prior to depletion reactions. ‘Supe’ refers to factor-depleted DREX supernatant from the immunoprecipitation (IP) reaction while ‘IP’ refers to protein fraction bound to the antibody-coupled beads. Approximately 20% of immunoprecipitated material was loaded. The target of IP is indicated above the blot. The bands corresponding to the detection target are indicated on the right side of the blot with the red asterisk marking a non-specific band. The position of the 135 kDa marker band is indicated to the left. c) 800-bp resolution Micro-C maps of the *eve* locus after Su(Hw) (top row) or Rad21 (bottom row) depletions. For each depletion condition; Left: Interaction maps representing balanced contact frequencies for control (below the diagonal) or depleted (above the diagonal) samples. Center: Differential bins (|log-fold change| >= 0.35 and p.value < 0.1) derived from statistical testing of indicated depletion versus control (with number of biological replicates used for testing denoted by N). Right: Difference maps representing log2-ratio of Depletion over Control contact frequencies. The dotted circle and arrow represent the Nhomie-Homie loop; called boundaries are denoted below. Scales for each heatmap type are shown to the right.

We then assembled chromatin using the factor-depleted extracts. As a readout for the quality of chromatin assembly, we quantified nucleosome array regularity from MNase-seq profiles with SDE scores at the previously described PNA classes. The resulting violin plots demonstrated that SDE scores were largely unchanged in chromatin assembled with DREX depleted of Su(Hw)-or Rad21, with the exception of a reduction observed at Su(Hw) motifs in Su(Hw)-depleted chromatin (Figure S7b). These results indicate that neither Su(Hw) nor Rad21 is required for robust chromatin assembly, although Su(Hw) directly contributes to positioning nucleosomes around its motif.

The Micro-C contact maps, differential bins and log-ratio difference maps for the different conditions (Figure 6c) showed that depletion of Su(Hw) significantly weakened the Nhomie-Homie loop and *eve* TAD, while depletion of Rad21 only led to very mild reduction of these long-range interactions. Some other, strong structures at different sites showed a similar trend. For example, the loop network on FlyFos029863 showed a drastic reduction in interaction frequencies in chromatin lacking Su(Hw) but not Rad21 (Figure S7c, top row). We also observed one instance (FlyFos019248) where Su(Hw) and Rad21 appeared to similarly weaken the loop connecting boundaries #27 and #30, while the loop connecting boundaries #29 and #30 was selectively perturbed only upon Su(Hw) depletion (Figure S7c, bottom row). Interestingly, loop #27 also shows a moderate enrichment of Rad21 binding (Figure 2).

In summary, these results argue against cohesin-mediated loop extrusion as the dominant mechanism of 3D chromatin structure formation in DREX-reconstituted chromatin, which instead likely relies on direct pairing-based mechanisms involving Su(Hw).

## Discussion

While in vivo studies have been instrumental in defining principles of genome organization, their complexity motivates complementary reconstitution approaches that allow mechanistic dissection of chromatin folding under controlled conditions. To this end, we built on our previously published chromatin assembly system using *Drosophila* syncytial blastoderm extracts (DREX) to reconstitute chromatin on long stretches of cloned *Drosophila* DNA and to probe the emergent folding of the nucleosome fiber using high-resolution Micro-C. This system, in combination with manipulation of DNA features, energy availability and protein abundance in the crude chromatin reconstitution extract, allowed us to interrogate mechanisms of *Drosophila* genome organization.

Pioneering studies have applied chromosome conformation capture-based approaches to reconstituted chromatin in an effort to derive fundamental principles of 3D genome organization. Fukai et al., applied in vitro Hi-C to long, synthetic nucleosome arrays to determine how polymer properties and histone acetylation modulate the folding of the synthetic fiber in the absence of proteins other than histones^4^. Oberbeckmann et al., reconstituted nucleosome arrays on cloned yeast DNA with genomic complexity and refined the nucleosome positions using nucleosome remodeling factors in combination with general DNA-binding factors^40^. Their Micro-C study showed that regularly phased nucleosome arrays punctuated by NFRs with bound transcription factors are sufficient to generate domain-like interaction patterns in the absence of loop extrusion or transcription. Together, these works demonstrate that aspects of genome folding can emerge from intrinsic chromatin properties.

While the abovementioned studies reconstituted nucleosomes from recombinant histones by salt gradient dialysis and thus generated a fully defined system of low protein complexity, our extract-based approach involves a high complexity of proteins of unknown or suspected functions and is therefore of more exploratory nature. While some properties of the chromatin reconstituted in extracts of syncytial blastoderm *Drosophila* embryos have been described, particularly those related to the remodeling, spacing and phasing of nucleosomes in the linear fiber^43^, the system has so far not been used to probe the emergent folding of chromatin. The earlier observation of abundant interactions of the Su(Hw) insulator complex with chromatinized genomic DNA^47^ motivated our efforts to explore the potential of the system to mount architectural features of chromatin that more closely resemble metazoan chromosomes. Micro-C profiling revealed long-range interactions of distant loci on the linear fiber (loops) as well as topologically associated domains (TADs) demarcated by pairs of boundary elements, with sizes ranging from ∼1-35 kb. Many of these chromatin loops are anchored at paired boundaries that are occupied by the Su(Hw) protein, as assessed by direct ChIP-seq experiments. We showed earlier that Su(Hw) associates with mod(mdg4), Map60 and Cul2 in DREX^47^. While our experiments establish that Su(Hw) is necessary for *eve* TAD formation, it is likely that the constitution of the *eve* boundaries requires other factors. The protein composition of DREX has been characterized to some degree^45^ and it is likely that other maternally deposited factors are present and contribute to chromatin folding. In line with this, assessing putative binding using published ChIP-seq datasets from cultured cells and embryos strongly argue that some of these factors, like promoter-related factor BEAF-32, contribute to functional boundaries in vitro. We did not find any evidence that Phaser, a prominent DNA binding protein known to create prominent arrays of phased nucleosomes in DREX-reconstituted chromatin^47^, engages in long-range interactions.

We also observed local punctuation of chromatin polymer interactions (‘insulation’) at individual sites, suggesting that insulation can arise from single boundary elements and does not require paired boundaries or well-defined domains.

Our comparison of folding patterns of chromatin reconstituted in extracts of syncytial blastoderm embryos with those of living embryos revealed striking differences. Notably, many in vivo TADs that appear at the onset of the major wave of zygotic genome activation (ZGA) (∼NC14) are not reconstituted. These structures may require developmental factors that are not yet present in the pre-ZGA embryos collected for DREX. Of note, the RNA polymerase II transcription machinery, which critically contributes to defining the embryonic chromosome architecture in *Drosophila*^56–58^, is inactive in DREX, which is supported by the fact that the commonly observed nucleosome phasing at promoters, a proxy for the assembly of preinitiation complexes, is not reconstituted in vitro^41,47^. Furthermore, only factors that are stored in the embryoplasm will be contained in DREX, since exact preparation involves removing the nuclei by centrifugation. Lowly abundant transcription factors, such as GAF and CLAMP, are therefore not present in DREX although they are maternally deposited^46^. We and others also showed that chromatin assembled by DREX lacks most abundant histone modifications, which limits the emergence of structures that are reinforced by chemical modifications, like Polycomb domains^44^. Missing factors may be supplemented in follow-up studies to provide instructive cues for further levels of chromatin organization.

We also observe several instances of Su(Hw)-mediated looping interactions that are not observed under any circumstance in embryos, despite the fact that the loop anchors are bound by Su(Hw) in vivo. Evidently, the in vitro system permits interactions that are suppressed in vivo by processes, structures or factors that are absent in DREX-reconstituted chromatin.

Alternatively, these differences may reflect variations in factor concentration, which can profoundly affect oligomerization/multimerization, rather than their absolute presence between the two systems. We cannot exclude that isolated regions cloned in our BACs and fosmids interact differently in vivo due to the chromosomal context of neighboring sequences that are missing in our constructs. Another possibility is that these interactions occur transiently in early embryos (which divide approximately every 8 minutes) making them technically challenging to capture. In contrast, the extended incubation in vitro allows these contacts to accumulate over time, increasing their probability of detection. A further source of discrepancy may arise from the fact that DREX is collected in a 90-minute time window from embryos that are heterogeneous with respect to developmental stage and asynchronous with respect to the cell cycle. The extract thus may well contain a mix of conflicting regulatory factors and signals, which collectively may promote non-physiological interactions. Of note, in vitro Micro-C data are often noisier than corresponding in vivo data. While this may be technical, it could also be a consequence of sampling a wider range of unconstrained and/or heterogeneous chromatin conformations in vitro, resulting in decreased signals and increased noise. In any case, resolving the reasons underlying the discrepancies in vitro and in vivo may turn out instructive and point to missing cues.

There is growing evidence that positioning of nucleosomes along the chromatin fiber can contribute to folding of the chromatin fiber^40,59–61^. Recent studies combining in vitro reconstitution, polymer simulations and in vivo perturbations have shown that intrinsic nucleosome array properties such as nucleosome spacing can influence short-range chromatin interactions. In human cells, perturbation of nucleosome density and phasing at CTCF sites through depletion of the histone chaperone FACT leads to weakening of TAD insulation despite largely preserved cohesin and CTCF occupancy^59^. These studies suggest that nucleosome positioning may contribute to fine-tuning chromatin boundaries. In vitro, Su(Hw) binding leads to robust nucleosome phasing on either side of the bound protein, similar to the situation of CTCF in human cells. However, nucleosome phasing around NFRs alone is insufficient to confer large-scale boundary activity since robust nucleosome phasing around Phaser binding sites do not serve as boundaries. Additional contextual features, such as NFR width in conjunction with cooperative architectural interactions, may be required for robust insulation. An efficient boundary may thus involve physical interactions between bound proteins as well as proper interdigitation of neighboring nucleosome arrays.

Our perturbation experiments were designed to distinguish between the two prevalent mechanisms, loop extrusion and direct boundary pairing, for formation of the *eve* TAD, for which we were able to obtain the most robust datasets. Although each of our approaches may leave room for alternative explanations, collectively the data are best explained by models in which the *eve* TAD is organized by direct pairing between protein complexes at Homie and Nhomie boundaries, in line with recent in vivo studies^11,51^. The immunodepletion of diagnostic factors from DREX suggests that the establishment of the *eve* TAD (and other in vitro loops) largely depends on Su(Hw) but not Rad21. These observations, in addition to arguing against Su(Hw) as a possible staller of extruding cohesin, establish Su(Hw) as a bona fide loop anchor. Once assembled, the TAD remains stable in the absence of ATP, similarly arguing against a mechanism involving continuous loop extrusion. Recent imaging studies showed that cohesin-driven extrusion is highly dynamic, with the lifetime of loops and TADs estimated to be in the scale of 5-30 mins^62,63^.

Cleavage of the *eve* locus with restriction enzymes allowed several interesting conclusions. First, interaction between the two boundary elements, Homie and Nhomie, are still observable if the DNA in-between the two anchors is cleaved and distance as well as orientation on the then linearized DNA molecule is massively altered. Further, we observe subtle residual interactions between Nhomie and Homie *in trans*, when the elements are on different DNA molecules. While we cannot rule out that these signals arise from a miniscule fraction of uncut molecules, Su(Hw) may well be able to bridge distant sites on different DNA molecules given its known role in pairing of homologous *Drosophila* chromosomes at specialized ‘button’ loci^64,65^.

Finally, we observe that interaction of *eve* gene sequences with either boundary is somewhat independent of contact between the loop anchors themselves. This argues for an instructive role of *eve* sequences in TAD formation rather than mere stochastic collisions of the fiber facilitated by a single loop extrusion event. In support of this, we observe possible binding of Zelda throughout the entire *eve* locus, in addition to its enrichment at the boundaries (Figure S1), raising the possibility that Zelda contributes to a cooperative chromatin environment that reinforces *eve*-centered interactions beyond the contribution of Su(Hw) alone.

While we propose with confidence that the eve TAD can form in vitro by direct pairing of Homie and Nhomie elements, this conclusion should not be generalized to all sites that may be reconstituted in vitro, nor to the in vivo situation. Loop extrusion may be generally compromised in the DREX system for unknown reasons or occur in specific genomic regions depending on local context. Given recent observations that many *Drosophila* boundaries are enriched for combinations of insulators and act in an orientation-dependent manner^66–69^, it is possible that some boundaries may impede the progress of loop extrusion by cohesin. Cohesin may also have a more general role; early studies which assessed the effect of dsRNA interference of Rad21 in cultured cells using Hi-C observed a small decrease in interactions in active regions of the genome (which are under-represented in DREX), partly phenocopying Pol II inhibition^57,70^. Given the ∼20-fold smaller genome of *Drosophila*, cohesin may play more specialized roles in facilitating local boundary interactions and gene regulatory contacts rather than serving as a dominant architectural ‘packaging’ factor, with extensive loop extrusion being less critical for overall chromatin organization.

Mindful of these shortcomings, we propose that in vitro reconstitution of complex, folded chromatin in extracts from early *Drosophila* embryos provides a novel approach for the investigation of chromatin organization that complements current in vivo analyses. The reconstitution system may be further developed by genome-wide assessment, detailed characterization of the DREX proteome, manipulation of cell cycle signaling and supplementation with transcription factors as well as chromatin organizers from later developmental stages. Most importantly, our study provides a first proof-of-principle that long-range chromatin folding can be reconstituted in vitro from soluble components that may stimulate follow up studies.

## Methods

*BAC/Fosmid pool preparation*. Bacterial stocks harboring BACs were acquired from BACPAC Resources, while fosmid bacterial stocks were generated as part of the FlyFos project^71^. Stocks were streaked onto LB agar plates (containing 12.5 µg/ml chloramphenicol for selection), incubated overnight at 37°C. Single colonies were picked and a 5 ml starter culture (LB medium and 12.5 µg/ml chloramphenicol) was inoculated overnight in bacterial shakers at 37°C, 130 rpm. Two ml of this culture were added to 100 ml LB medium supplemented with 12.5 µg/ml chloramphenicol and grown to an OD600 of ∼ 0.2. Arabinose was added to 0.1% w/v (f.c.) to induce replication of the BAC/fosmid and the culture was grown for another 5 hr at 37°C. Cells were harvested by centrifuging (4000 g, 20 mins) and pellets were kept frozen at −70°C until purification. BAC/fosmids were purified with a Machery-Nagel Nucleobond Xtra Midiprep kit. Washed DNA pellets were dissolved in TE, DNA concentrations were measured using Nanodrop spectrophotometer and the integrity of the DNA was assessed using the Agilent Tapestation gDNA system. In order to minimize accidental shearing/linearization of the circular DNA, usage of narrow pipette tips and freeze-thaw cycles were avoided. For all experiments, equimolar pools of final concentration of ∼200 ng/µL were prepared by accounting for the different lengths of the constructs.

*Restriction endonuclease cleavage*. Restriction enzymes were acquired from New England Biolabs and used according to manufacturer’s recommendations to digest the BAC/fosmid pools (at a ratio of 2 units enzyme/1 µg DNA pool). Following the digestion, DNA was precipitated overnight at −20°C directly by addition of 1 volume 7.5 M ammonium acetate, 2.5 volumes of 100% ethanol and 2 µL of 20 µg/µL glycogen. The pellet was washed twice with 70% ethanol, reconstituted in TE and concentration was measured prior to use in DREX assembly.

*Drosophila embryo extract (DREX) preparation*. DREX was prepared according to Eggers et al^46^. Briefly, embryos were collected from Oregon-R fly population cages repeatedly in 90-minute intervals^41^. To minimize contamination from later embryos, collected embryos were accumulated on ice to arrest further development. 50 ml of settled embryos were dechorionated in 200 ml embryo wash buffer (0.7% NaCl, 0.04% Triton-X100 in water) and 60 ml 13% sodium hypochlorite (VWR Chemicals) for 3.5 min at room temperature (RT) with stirring using magnetic stirrers. Embryos were rinsed for 5 min on a sieve with cold water and transferred into a glass cylinder with embryo wash buffer. Floating debris was aspirated off and settled embryos were subsequently washed first in 0.7% NaCl solution and then in cold EX10 buffer (10 mM HEPES pH 7.6, 10 mM KCl, 1.5 mM MgCl2, 0.5 mM EGTA and 10% v/v glycerol) freshly supplemented with 1 mM DTT and protease inhibitors (0.2 mM PMSF, 1 mM Leupeptin, 1 mM Aprotinin, 1 mM Pepstatin). Embryos were settled in a Homogenplus 50 ml homogenizer (Schuett-Biotec), the supernatant was aspirated and the embryos were homogenized with one stroke at 3000 rpm and 10 strokes at 1500 rpm. The MgCl_2_ concentration of the homogenate was adjusted to 5 mM (final concentration) and centrifuged for 10 min at 13,000 g at 4°C. This step results in separation of multiple phases: a white lipid layer on top, debris at the bottom with a clear solution in the middle. The middle layer was further centrifuged for 2 h at 245,000 g at 4°C. The clear extract was collected with a syringe, leaving the lipid layer and pellet behind. Extracts were stored in 200 µl aliquots - 70°C after flash freezing in liquid N2. Extracts were only thawed once before use.

*Chromatin assembly*. Chromatin was assembled by addition of 100 µL DREX to 1 µg of BAC/Fosmid pool along with 15 µL 10x McNAP buffer (30 mM MgCl2, 10 mM DTT, 300 mM creatine phosphate, 30 mM ATP, 10 µg/ml creatine phosphate kinase (CPK in 100 mM imidazole pH 6.6)) and 0.5 µL of 1 mM ZnCl_2_. The reaction volume was adjusted to total 150 µL using EX50 buffer (10 mM HEPES ph 7.6, 50 mM NaCl, 1.5 mM MgCl_2_, 10% (v/v) glycerol). The assembly reaction was vortexed for 5 seconds on low settings and incubated for 16-18 hrs at 26°C 300 rpm on a tabletop Thermomixer.

*Immunodepletion*. To deplete proteins of interest from DREX prior to assembly, we adopted an antibody-mediated immunoprecipitation strategy. Per target, 30 µL of prewashed (blocked) protein-G magnetic Dynabeads (Thermo Fisher Scientific) was precoupled to 7 µL antibody sera (polyclonal rabbit antibodies raised against the full length Su(Hw) protein from Dr. Carla Margulies, LMU Munich; polyclonal rabbit antibodies raised against amino acids 482–702 of Rad21 from Prof. Dr. Claudio E. Sunkel, University of Porto) diluted in 500 µL cold EX50 buffer supplemented with Roche Protease Inhibitor Cocktail. A sample containing an equivalent amount of a non-specific rabbit anti-IgG antibody (Cell Signaling Technology, #2729) was included as a control to account for bead ‘stickiness’. The precoupling was performed for 90 mins at 4°C on a rotating wheel. Bead-antibody complexes were blocked for 30 mins with 3% (w/v) Bovine Serum Albumin (BSA) solution in EX50 at RT and quickly washed in cold EX50 before the next step. The blocked bead-antibody complexes were incubated with 140 µL of DREX in 1.5 ml low binding tubes. Tubes were placed on ice and gently flicked/pipetted periodically for 3 hours. Subsequently, beads were magnetically separated from the supernatant. To assess for depletion efficiency by Western blotting, 30 µL of the supernatant was mixed with 7.5 µL 5x Laemmli Buffer (along with an equivalent volume of unprocessed DREX as Input) while the beads were directly resuspended in 55 uL 1x Laemmli Buffer. All samples boiled at 95°C for 5 mins and analysed by SDS gel electrophoresis.

*Western blotting.* Samples were electrophoresed on SDS ServaGel TGPrimer 8% gels for 1.5–2 h at 180 V. Proteins were transferred to Amersham™ Protran™ 0.45-μm nitrocellulose blotting membrane for 1.5 h at 300–400 mA in either transfer buffer (20% MeOH, 25 mM Tris, 192 mM glycine). Membranes were blocked with 3% BSA for 1 h at RT. The membrane was incubated with primary antibody (diluted 1:1000) overnight at 4°C in 3% BSA, washed thrice with PBS-T (PBS/0.1% Tween 20) and incubated with secondary antibody in PBS-T for 1 h at RT. Images were acquired and quantified using the LICOR Odyssey CLx. For all quantifications, we verified that the signal is linear in the range of loading used based on serial dilutions.

*ATP measurement and manipulation*. ATP quantifications were performed using the CellTiter-Glo Luminescence Assay Kit (Promega) and measured on the Promax Navigator instrument.

The reagent was determined to work approximately linearly in the range of ATP concentrations between 10 µM and 10 nM based on serial dilutions. For all DREX measurements, 1 µL of DREX assembly reaction was diluted in 499 µL EX50. 50 µL of this dilution was mixed with 50 µL of the reagent, pipetted onto an opaque bottom 96-well plate (along with appropriate standards and background wells) and incubated for 10 mins prior to measurement. All measurements were performed as triplicates to account for technical variability.

To rapidly deplete ATP from the assembly reaction, Apyrase (NEB) was diluted 1:5 in the corresponding buffer and 5-7.5 µL of this diluted apyrase was added to 150 µL of the chromatin assembly reaction. As ATP depletion results in a charge imbalance and precipitation of chromatin, 7.5 µL of 100 mM ATP-γS stock solution was added to the ∼160 µL of ATP-depleted DREX chromatin samples.

*In vitro Micro-C and MNase-seq*. In vitro Micro-C was adapted from Oberbeckmann et al^40^. After overnight assembly, the chromatin was double-crosslinked by addition of 3 mM (f.c.) DSG for 20 mins and formaldehyde for 10 mins (0.2% f.c.) at 26°C. Excess crosslinkers were quenched by addition of 40 µL 5x Quenching Solution (250 mM Tris, 400 mM aspartic acid, 100 mM lysine, pH 8) to the assembly reaction. Samples were then processed for MNase digestion. MNase stock solutions were prepared by dissolving 500 U MNase (Sigma) in 850 µL EX50 buffer. 265 µL of digestion solution (259 µL EX50, 1.325 µL 1 M CaCl_2_, 4.5 µL of stock MNase) was added to 200 µL of crosslinked chromatin resulting in digested fragments with a mono-:di- :trinucleosome ratio of 50:25:15 (typically 75 sec for most conditions). Based on our optimization experiments, in vitro chromatin has a tendency to be overdigested by MNase (resulting in inefficient ligation) with prolonged digestion times. After digestion, 25 µL of 0.5 M EDTA was added to the reaction to inhibit MNase activity.

Digested chromatin was then purified to eliminate small subnucleosomal fragments (which may interfere with ligation). To do this, chromatin was diluted two-fold in EX50 buffer and concentrated on 50 kDa Amicon centrifugal filters (2000 g x 3 mins) for a total of three times. 20% of purified chromatin was taken as digestion control (MNase-seq samples) while the rest was used to set up the Micro-C proximity ligation, in which the DNA end repair, phosphorylation and ligation steps are performed in a single tube. Chromatin was resuspended in T4 Ligation Buffer (NEB) and was supplemented with final concentrations of 0.35 mM dNTP, 2.5 mM EGTA, 10 Cohesive End Units (CEU)/µL T4 Ligase (NEB), 0.025 U/µL Klenow (NEB) and 0.05 U/µL Polynucleotide Kinase (NEB). The ligation reaction was incubated for 2 hrs at 37°C and 8 hrs at 20°C on a Eppendorf Thermomixer set at 300 rpm. After ligation, RNAase treatment was performed for both ligation and digestion control samples for 30 mins at 37°C after which decrosslinking was performed at 65°C for 3 hrs in the presence of Proteinase K (0.6 µg/µL f.c.) and SDS (0.2% f.c.) with vigorous shaking. Subsequently, DNA fragments were purified by phenol-chloroform extraction and quantified using Qubit reagent. Digestion and ligation efficiencies were assessed using the HSD5000 Agilent Tapestation system.

Sequencing libraries for Micro-C samples were generated as follows. The volumes of ligation samples were adjusted to 50 uL using TE and unligated mononucleosomes were eliminated by size selection by the addition of 45 µL AMPure XP beads. Size selected DNA was eluted in 140 µL and sheared uniformly to ∼200 bp in 130 µL Covaris SnapCap microTubes on the Covaris S220 instrument using the following settings: peak incident power of 175, duty factor 10%, 200 cycles per burst for 250 sec at 4°C. Sheared DNA was purified by addition of 234 µL of AMPure XP beads, products eluted in 52 µL TE and quantified using Qubit (resulting in typical yields of 50-200 ng). These samples were then processed using NEBNExt Ultra II library preparation kit following manufacturer’s recommendations until the PCR amplification step. Prior to PCR amplification, the adaptor ligated samples were split into two technical replicates to minimize amplification biases. PCR amplification was performed on the first batch using Agilent Herculase II reagents along with NEB Dual indices for Illumina using the following PCR settings.

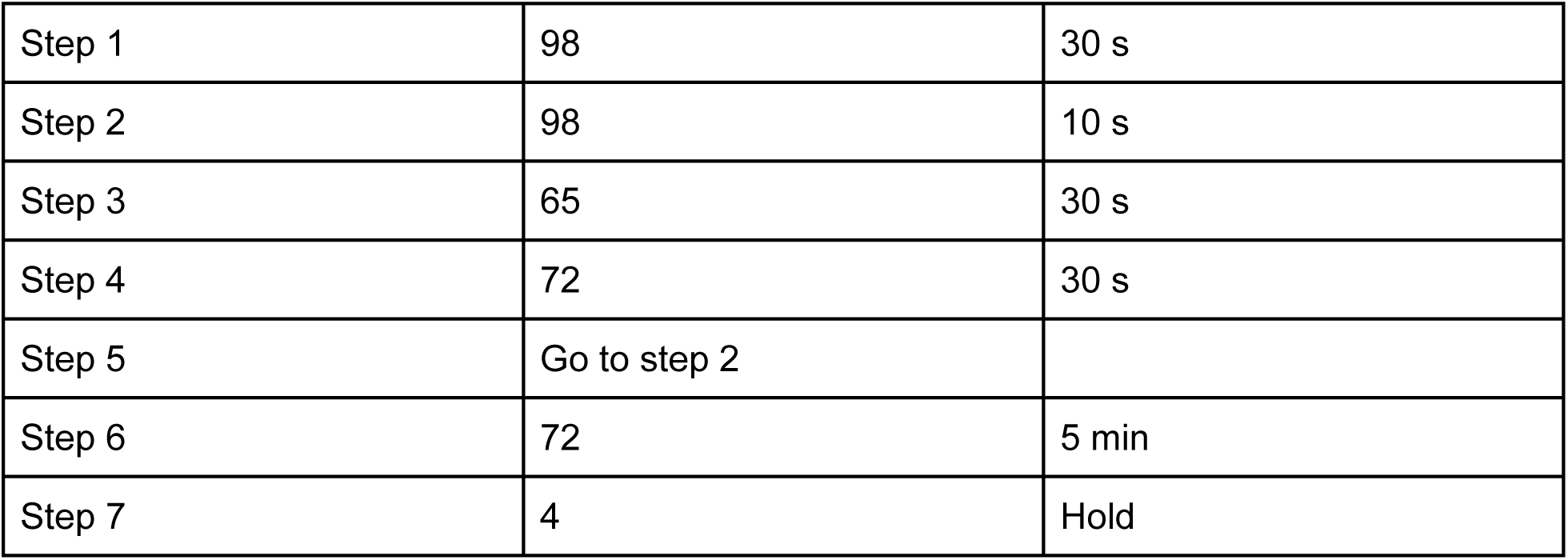

1 µL of the PCR reaction was run on the HSD1000 Tapestation system to verify the quality of PCR amplification and to assess the number of required amplification cycles (typically 4-6). The second batch was then processed identically, pooled with the first batch and size selected two consecutive times using AMPure XP beads to eliminate primer dimers. Libraries were sequenced on an Illumina NextSeq 1000 instrument using 60 bp x 2 paired-end settings to yield roughly 25 million reads per sample per replicate.

For MNase-seq, 10 ng of digestion control samples were processed using the NEBNext Ultra II kit following manufacturer’s protocol. Generated libraries were size selected for mononucleosomes using AMPure XP beads prior to sequencing to a depth of 3 million reads per sample per replicate.

*In vitro MNase ChIP-seq.* Assembled chromatin was crosslinked with formaldehyde (0.1% f.c.) for 5 mins at 26°C and subsequently quenched using fresh glycine solution (150 mM f.c.). Crosslinked chromatin was digested using Micro-C digestion solution (see above) for 3 mins and the MNase was inactivated by addition of EDTA (25 mM f.c.). Digested chromatin was precleared using 20 µL equal mixture of proteinA/G Sepharose Fast-Flow beads (cat #) for 1 hr at 4°C on a rotating wheel. In parallel, 20 µL mixture of proteinA/G beads were precoupled to 1 µL of the Su(Hw) antibody for 3 hrs at 4°C on a rotating wheel. A small amount of precleared chromatin was taken as input and the rest was used for overnight IP reaction by addition of antibody-bead complexes. RNAse treatment, heat decrosslinking using Proteinase K/SDS and purification using Phenol Chloroform was performed similar to the Micro-C samples. Sequencing libraries were generated similar to MNase-seq but without the mononucleosome size selection to retain small fragments often associated with transcription factor binding.

## Data analysis

*Fosmid selection and custom genome assembly*. To select a subset of interesting fosmids from a list of 742 fosmids tiling the X-chromosome, we first calculated a cumulative score representing the number of occurrences of features like the Su(Hw) motif, Zelda ChIP-seq peaks, Hi-C boundaries defined in *Drosophila* S2 cell line etc. High-scoring fosmids were selected for purification as detailed above. A custom genome was assembled based on the dm6 coordinates of the inserts of the 24 fosmids (FlyFos prefix) and the 4 BACs (MG prefix) spanning the *eve* locus. Each construct was named and identified throughout the study as follows: prefix followed by unique numerical identifier and original dm6 coordinates. This custom genome was used throughout the study for alignment of sequencing data.

*Micro-C data processing and analyses*. Raw reads were processed using the nf-core/hic pipeline with the following parameters ‘--dnase --min_cis_distance 150’ for HiC-Pro processing and ‘--mad-max 12 --min-nnz 3 --min-count 1’ for cooler balancing. 17 independent control Micro-C experiments were pooled together to generate a high-complexity reference dataset used for subsequent analyses. Generated .mcool files (containing contact matrices of 200, 400, 800, 1000, 1600, 3200 bp resolutions) were used for visualization and analyses with the Bioconductor workflow OHCA^72^ in R version 4.3. A list of blacklist regions was defined based on regions of low Micro-C coverage in the pooled dataset. Further, overlapping regions between fosmids/BACs as well as the ends were also included in the blacklist. For all feature-based analyses in this study (for instance, boundary calling, nucleosome phasing around motifs etc.), we filtered out features within 1 kb of any blacklisted region. Diamond insulation profiles were calculated using 800,1600 and 3200bp sliding windows using the 200 bp resolution contact map generated from pooled control samples. As the profiles were highly concordant, the 1600 bp-based insulation track was used for subsequent analyses. Local minima in insulation scores were called as boundaries and were subsequently filtered for a score of equal to or lower than - 0.5 to retain high-confidence, strong boundaries. For analyses of the experiments involving perturbations of DNA sequences, ATP and DREX content, we adopted a statistical approach using the Bioconductor package multiHiCCompare, which allows joint normalization and difference detection of HiC-style data^73^. Briefly, .mcool files for individual replicates were converted into hicexp matrices containing interaction raw counts (with A.min parameter set to 2) along with information on sample groups (for instance, based on the restriction enzyme used) and experimental batch. Joint normalization was performed using cyclic loess and difference detection was performed using GLMs (quasi-likelihood model) with sample groups and experimental batches as covariates. Statistically-differing bins (abs(log2(Fold Change)) >= 0.35 and p.value <= 0.1) were visualized adjacent to standard contact maps from pooled data. Division-based difference heatmaps were generated by dividing contact maps from treatment samples by control samples (with pseudocount 0.01). The resulting heatmap was smoothened by despeckling with the focal size parameter to 1.

*MNase-seq data processing and analyses*. MNase-seq data processing was performed based on Baldi et al.^47^. Briefly, raw reads were aligned to the custom genome using bowtie2^74^ with parameters “-p 20 -X 250 --no-discordant --no-mixed --no-unal”. Subsequent steps were performed with R version 4.0, primarily relying on Bioconductor packages (e.g., GenomicRanges, GenomicAlignments) for genomic analyses and the tidyverse ecosystem for data wrangling and visualization. Dyad coverage vectors were obtained by size-selecting fragments of length > = 120 and ≤ 200 bp and resizing their length to 50 bp fixed at the fragment center. These resized fragments were piled up to determine coverage while adjusting for copy number differences between the different constructs within the pool. Nucleosome phasing was assessed by scanning dyad coverage profiles with a sliding-window spectral density estimation (SDE) approach as used in Baldi et al.,. For each 1024 bp window, a periodogram was computed and spectral density at the nucleosome repeat length (179 bp; estimated based on autocorrelation of MNase-seq coverage) was extracted and z-score normalized for each construct. Phased Nucleosome Arrays (PNAs) were defined as regions where spectral density z-score >= 2 and are at least 550 bp. PNAs were further stratified into 3 categories: Su(Hw)-type, Phaser-type and other, based on overlap (or lack of) with previously described sequence motifs. 44 randomly generated centers (weighted based on construct size) served as controls. To visualize MNase-seq coverage around 3-5 kb windows of different sites of interest as composite plots or heatmaps, the Bioconductor package HelpersforChIPseq was used. For correlating SDE scores with insulation, the windows were restricted to 500 bp around motifs to calculate mean feature values. Restriction digestion sites were predicted using jvarkit^75^.

*MNase ChIP-seq data processing and analyses*. MNase ChIP-seq data processing was performed based on Jayakrishnan et al.^76^. Briefly, raw reads were mapped to the custom genome using bowtie2^74^ with parameters “-p 12 --end-to-end --very-sensitive --no-unal --no-mixed --no-discordant -I 10 -X 500”. Subsequently, HOMER^77^ was used to generate library-size and input-normalized coverage tracks which were visualized on Integrative Genomics Viewer (IGV)^78^.

*Published ChIP-seq data processing and analyses.* Full list of external datasets used in this study is provided in the data availability section. Coordinates of ChIP-seq datasets aligned to dm3 were converted into dm6 using liftOver^79^. All dm6 datasets were then coordinate-transformed to the custom genome based on starts/ends of the BAC/fosmid constructs. To facilitate uniform comparison of ChIP-seq datasets originating from different sources, we first resized the coverages into 200 bp bins and then performed a log quantile normalization as previously described^24^. To analyze the enrichment of different factors at Micro-C boundaries, normalized coverages were loaded and average signals within the 1kb window around the boundary centers were calculated and visualized as a heatmap using the Bioconductor package ComplexHeatmap^80^.

## Data availability

The Micro-C, MNase-seq and ChIP-seq datasets generated for the current study are available from the Gene Expression Omnibus (GEO) as a SuperSeries under accession number GSE317985 (reviewer token : edmzyywapdklfcf). The following external datasets were used in this study:

| Dataset (Identifier) | Source |
| --- | --- |
| Zelda ChIP-seq 0-2hr embryos (GSE152598) | Duan et al.,<sup>81</sup> |
| BEAF-32 ChIP-seq Kc cells (GSM1535963) | Li et al.,<sup>70</sup> |
| Cap-H2 ChIP-seq Kc cells (GSM1535967) | Li et al.,<sup>70</sup> |
| CBP ChIP-seq Kc cells (GSM1535971) | Li et al.,<sup>70</sup> |
| Chromator ChIP-seq Kc cells (GSM1535976) | Li et al.,<sup>70</sup> |
| CP190 ChIP-seq Kc cells (GSM1535980) | Li et al.,<sup>70</sup> |
| CTCF ChIP-seq Kc cells (GSM1535983) | Li et al.,<sup>70</sup> |
| DREF ChIP-seq Kc cells (GSM1535985) | Li et al.,<sup>70</sup> |
| Fs1h-L ChIP-seq Kc cells (GSM1535988) | Li et al.,<sup>70</sup> |
| L3mbt ChIP-seq Kc cells (GSM1536002) | Li et al.,<sup>70</sup> |
| Rad21 ChIP-seq Kc cells (GSM1536010) | Li et al.,<sup>70</sup> |
| RNAPII ChIP-seq Kc cells (GSM1536014) | Li et al.,<sup>70</sup> |
| TFIIIC ChIP-seq Kc cells (GSM1536020) | Li et al.,<sup>70</sup> |
| Z4 ChIP-seq Kc cells (GSM1536022) | Li et al.,<sup>70</sup> |
| Ser5-Phospho Pol II ChIP-seq NC12 embryos (GSM1536376) | Blythe et al.,<sup>82</sup> |
| Ser5-Phospho Pol II ChIP-seq NC14L embryos (GSM1536386) | Blythe et al.,<sup>82</sup> |
| Pita ChIP-seq 0-12 hr embryos (GSM2042225) | Zolotarev et al.,<sup>83</sup> |
| ZIPIC ChIP-seq 0-12 hr embryos (GSM2042226) | Zolotarev et al.,<sup>83</sup> |
| Zw5 ChIP-seq 0-12 hr embryos (GSM2042227) | Zolotarev et al.,<sup>83</sup> |
| ATAC-seq NC11 embryos (GSM2219678) | Blythe et al.,<sup>84</sup> |
| ATAC-seq NC12 embryos (GSM2219696) | Blythe et al.,<sup>84</sup> |
| ATAC-seq NC13 embryos (GSM2219700) | Blythe et al.,<sup>84</sup> |
| Su(Hw) ChIP-seq Kc cells (GSM5945681) | Chen et al.,<sup>85</sup> |
| Su(Hw) ChIP-seq 0-24 hr embryos (GSM8109740) | modENCODE<sup>86</sup> |
| Hi-C TAD boundaries Kc cells (GSE97965) | Ramirez et al., <sup>29</sup> |
| Hi-C TAD boundaries S2 cells (GSE101317) | Wang et al., <sup>87</sup> |
| Low-input Micro-C NC9-14 embryos (EBI ArrayExpress ID: E-MTAB-14477) | Maziak et al., <sup>26</sup> |

Source data are provided with this paper.

## Acknowledgements

Bacterial stocks carrying X-chromosomal fosmids were a kind gift from Dr. P. Tomancak (Central European Institute of Technology, Brno, Czech Republic). We thank Dr. C. Sunkel (University of Porto, Portugal) for the rabbit anti-Rad21 antibody. We are grateful to E. Lei (National Institutes of Health, USA) for sharing the guinea pig anti-Su(Hw) antibody for orthogonal verification of our ChIP-seq datasets generated using the rabbit-Su(Hw) antibody. ATP-γS was a kind gift from Dr. K-P. Hopfner (Gene Center, LMU Munich, Germany). We thank all members of the Becker lab for helpful discussions. We thank Dr. T. Straub from the BMC Bioinformatics Unit for access to the computational cluster and bioinformatics advice. We thank S. Krebs of LMU LAFUGA for Illumina sequencing. We thank Y. Zhu from Oudelaar lab for help with Micro-C data processing. M.J. received support from the International Max-Planck Research School (IMPRS) “Molecules of Life” and the Integrated Research Training Group of the DFG-funded Collaborative Research Center “Chromatin Dynamics” during the project duration. Schematics in the manuscript were created on BioRender.com.

## Contributions

M.J. designed and performed all experiments and data analyses. G.K. prepared DREX. A.C-S. assisted in optimization of chromatin assemblies and preparation of sequencing libraries. M.K. and A.M.O. provided advice for optimization of the in vitro Micro-C protocol as well as data processing. N.M. and J.M.V. provided early access to Pico-C data and contributed to in vivo analyses. S.Z., C.E.M. and M.C.G. provided key reagents and advice. M.J. and P.B.B. wrote the manuscript with inputs from all authors. P.B.B. acquired funding and supervised the study.

## Ethics declaration

Competing interests:

The authors declare no competing interests.

## Code availability

The custom code used for Micro-C, MNase-seq and ChIP-seq data analysis is available on Zenodo (https://doi.org/10.5281/zenodo.18846930).

## Supplementary data

**Supplementary Figure S1.**
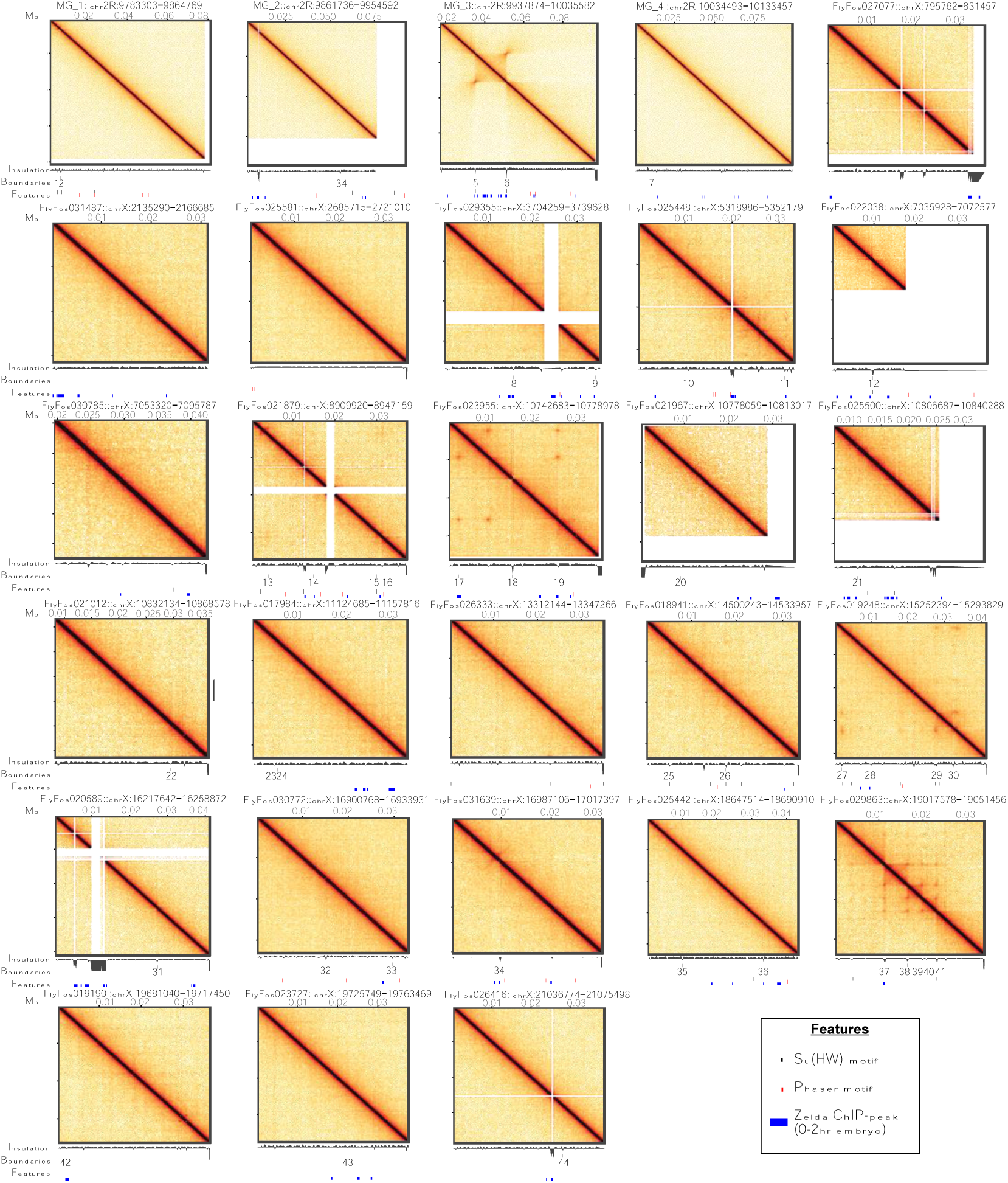
**Complex folding structures are observed in a subset of BACs/fosmids** 200-bp resolution Micro-C contact maps for all 28 constructs used in this study. The BAC/fosmid identifier and distance from insert start is indicated above each heatmap. All heatmaps were scaled uniformly as in Figure 1d. Diamond insulation scores as well as called boundaries with index numbers are indicated below the heatmap for reference. Su(Hw) and Phaser motifs as well as published 0-2 hr embryo Zelda ChIP-seq peaks (coloured based on the shared legend at the bottom) serve as an interpretive guide.

**Supplementary Figure S2.**
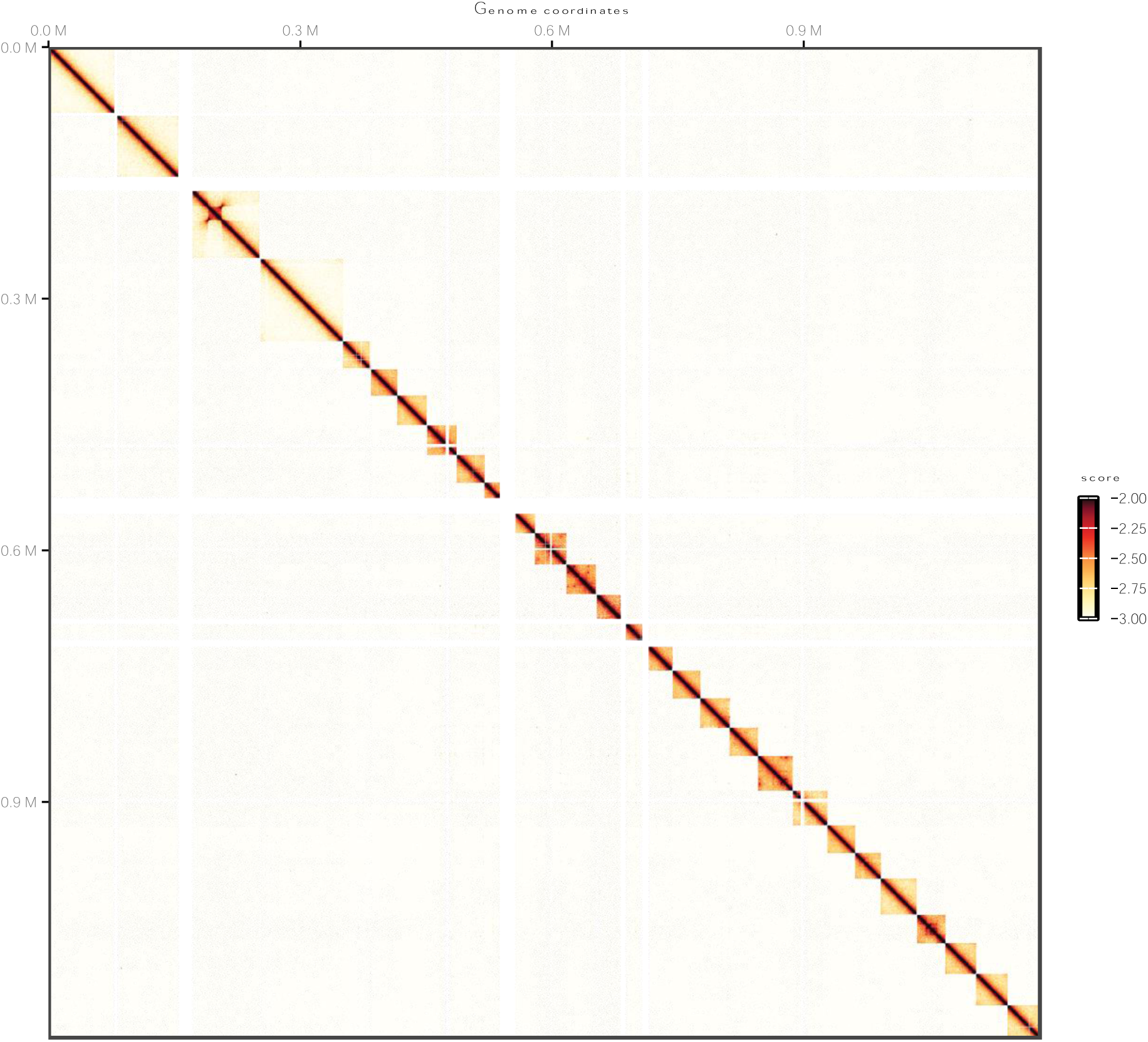
**No prominent *trans* interactions observed in the chromatinized BAC/fosmid pool** 400-bp resolution Micro-C contact map representing the absence of inter-construct interactions. Heatmap scale is provided on the right.

**Supplementary Figure S3.**
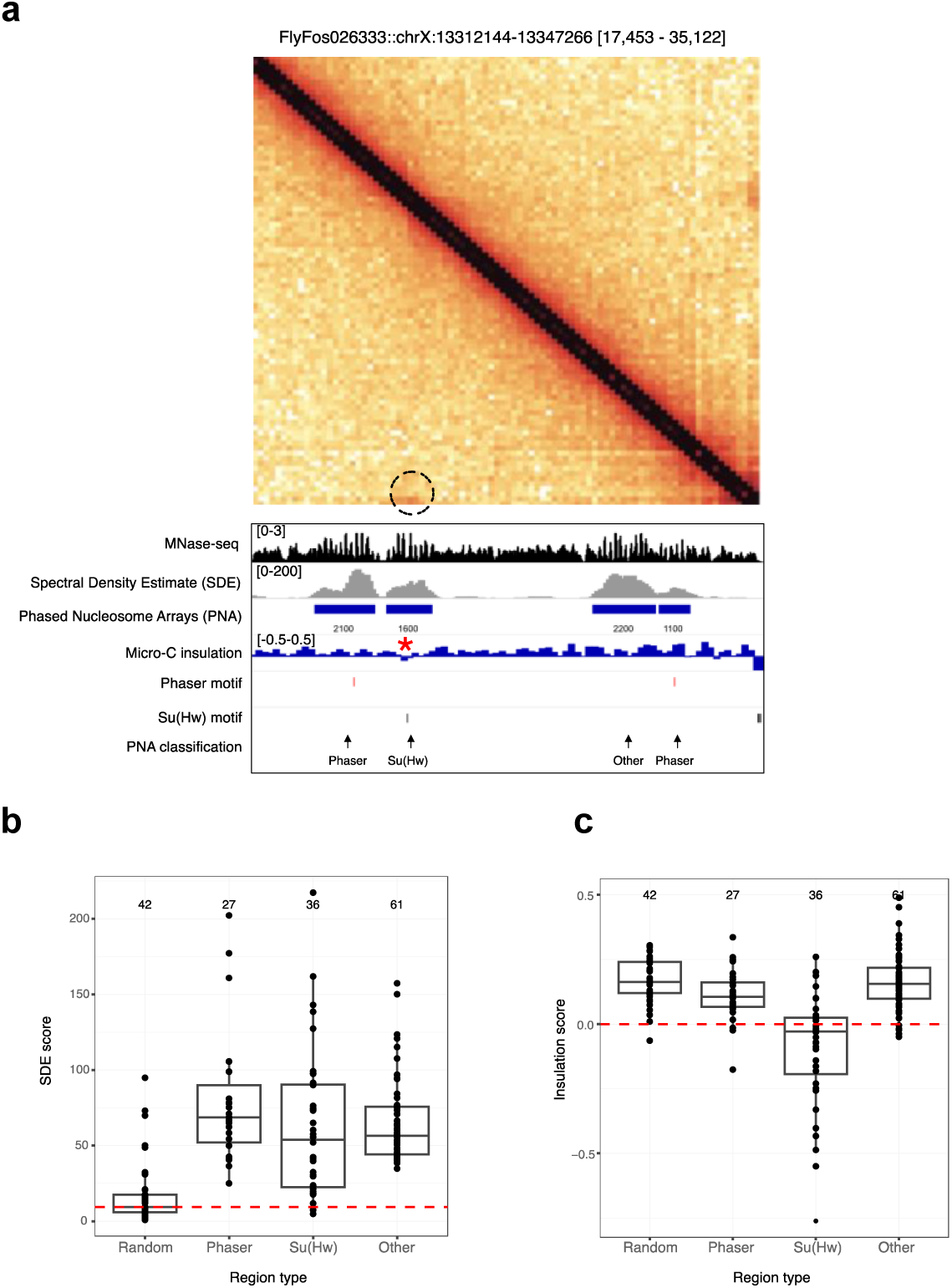
**Nucleosome phasing correlates poorly with Micro-C insulation** a) Top: 200-bp resolution Micro-C contact map for FlyFos026333. Subset region is indicated in square brackets. Dotted circle highlights a looping interaction. Bottom: MNase-seq profile, periodicity Spectral Density Estimate (SDE) scores along with extracted Phased Nucleosome Arrays (PNAs) and Micro-C insulation profiles for the same genomic region as shown in Micro-C contact map. The numbers below the PNAs indicate the PNA lengths in base pairs. The red asterisk on the diamond insulation track highlights the region with local minimum (-0.42) in insulation score. Phaser and Su(Hw) motifs along with PNA classification (based on overlap with motifs) are marked for reference. b) Boxplots representing the distribution of average SDE scores in 0.5 kb windows around centers of PNAs grouped by type. Random regions serve as a baseline reference. The median SDE score of random regions is represented by the red dotted line. The number of data points in each group is indicated above each box. c) Boxplots representing distribution of average Micro-C diamond insulation scores in 0.5 kb windows around centers of PNAs grouped by type as in (b). The red dotted line demarcates insulating (score < 0) and non-insulating (score >= 0) regions. The number of data points in each group is indicated above each box. Numerical values are provided in source data.

**Supplementary Figure S4.**
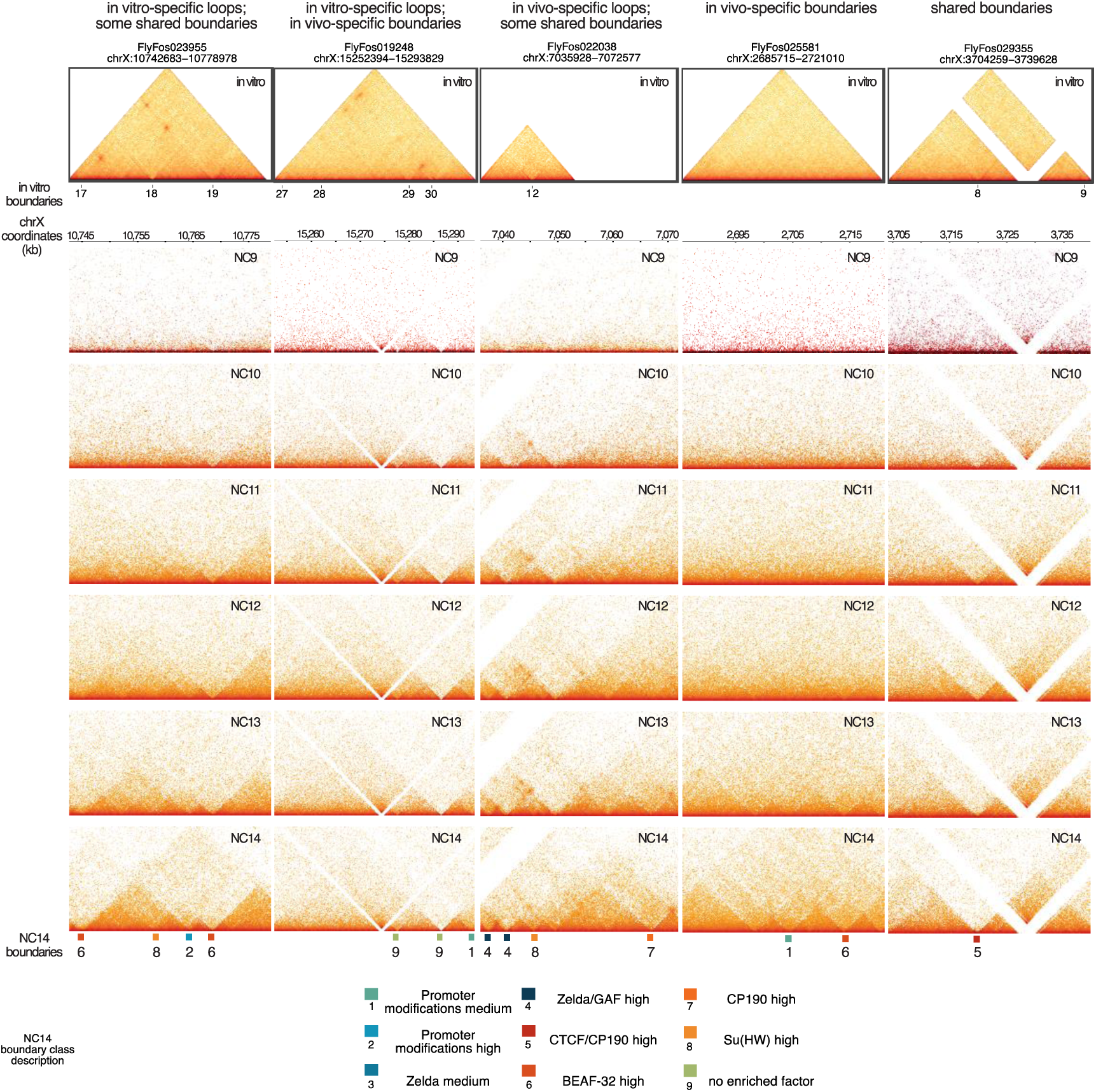
**Additional examples for discrepancies between in vivo and in vitro 3D chromatin folding patterns** Top: 200-bp resolution horizontal Micro-C contact maps for selected DREX-chromatinized regions. A brief description of notable patterns based on correlation with in vivo data is indicated at the top. Region identifier is indicated above. Identified in vitro boundaries along with reference index are shown below the contact heatmap. Below: 125-bp resolution Pico-C contact maps from developing Drosophila embryos for in vitro matched genomic regions. Developmental stages indicated by NC9-14 are shown in the top-right corner of the contact map. Chromosomal coordinates (dm6) are indicated above. Positions of boundaries identified in NC14 embryos as well as boundary classes are shown below NC14 contact maps. Brief descriptions of in vivo boundary classes are shown as a legend at the bottom.

**Supplementary Figure S5.**
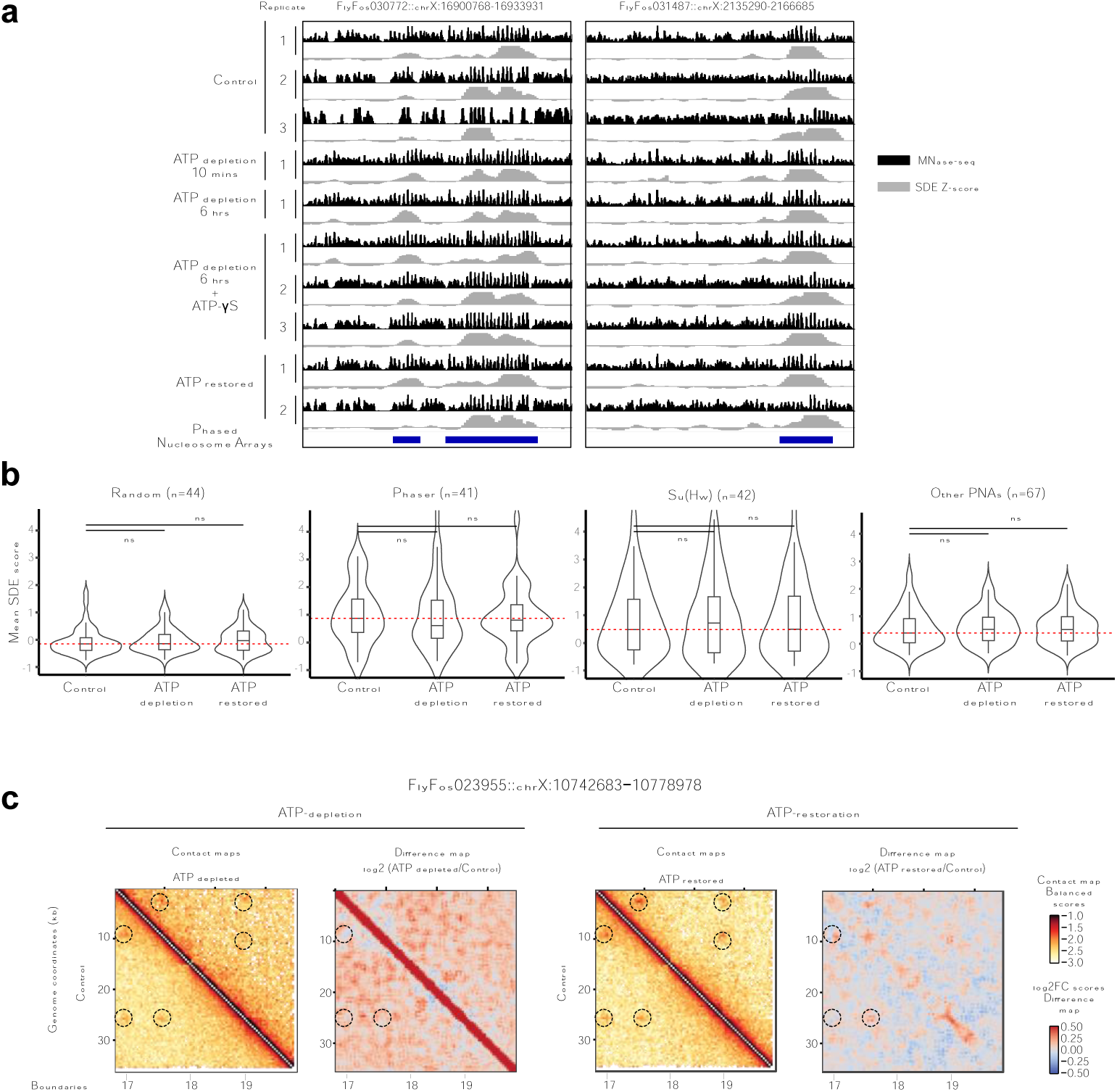
**Effects of ATP levels on nucleosome positioning and 3D genome structures** a) MNase-seq profiles (black) and associated SDE score tracks (gray) for individual replicates from ATP perturbation experiments for two representative regions. The construct identifier for regions of interest is shown above. Replicate number as well as type of perturbation is indicated on the left. The positions of Phased Nucleosome Arrays serve as a reference. b) Violin plots indicating distributions of replicate-averaged, mean periodicity SDE scores in 1-kb windows around centers of PNAs for Control, ATP-depleted and ATP-restored conditions grouped by PNA classes. Random PNAs serve as a baseline reference and an internal negative control. The red dotted line indicates the median SDE-score for the corresponding control sample. Two-sided, paired Wilcoxon-signed rank tests were performed per PNA class to compare each treatment condition against the control and the resulting p-values corrected for multiple comparisons using the Benjamini-Hochberg method. The significance values are indicated as ‘ns’ (non-significant), ‘*’ (p-adjusted < 0.05) or ‘**’ (p-adjusted < 0.01). Adjusted p-values for all comparisons are provided in associated Source Data. c) 400-bp resolution Micro-C maps of FlyFos023955 characterizing changes in contact patterns upon either ATP depletion (left) or ATP restoration (right). For each treatment condition, interaction maps representing balanced contact frequencies for control (below the diagonal) or treated (above the diagonal) samples are on the left while difference maps representing log2-ratio of Treatment over Control contact frequencies are on the right. The dotted circles represent loops and called boundaries are denoted below. Scales for each heatmap type are shown on the right.

**Supplementary Figure S6.**
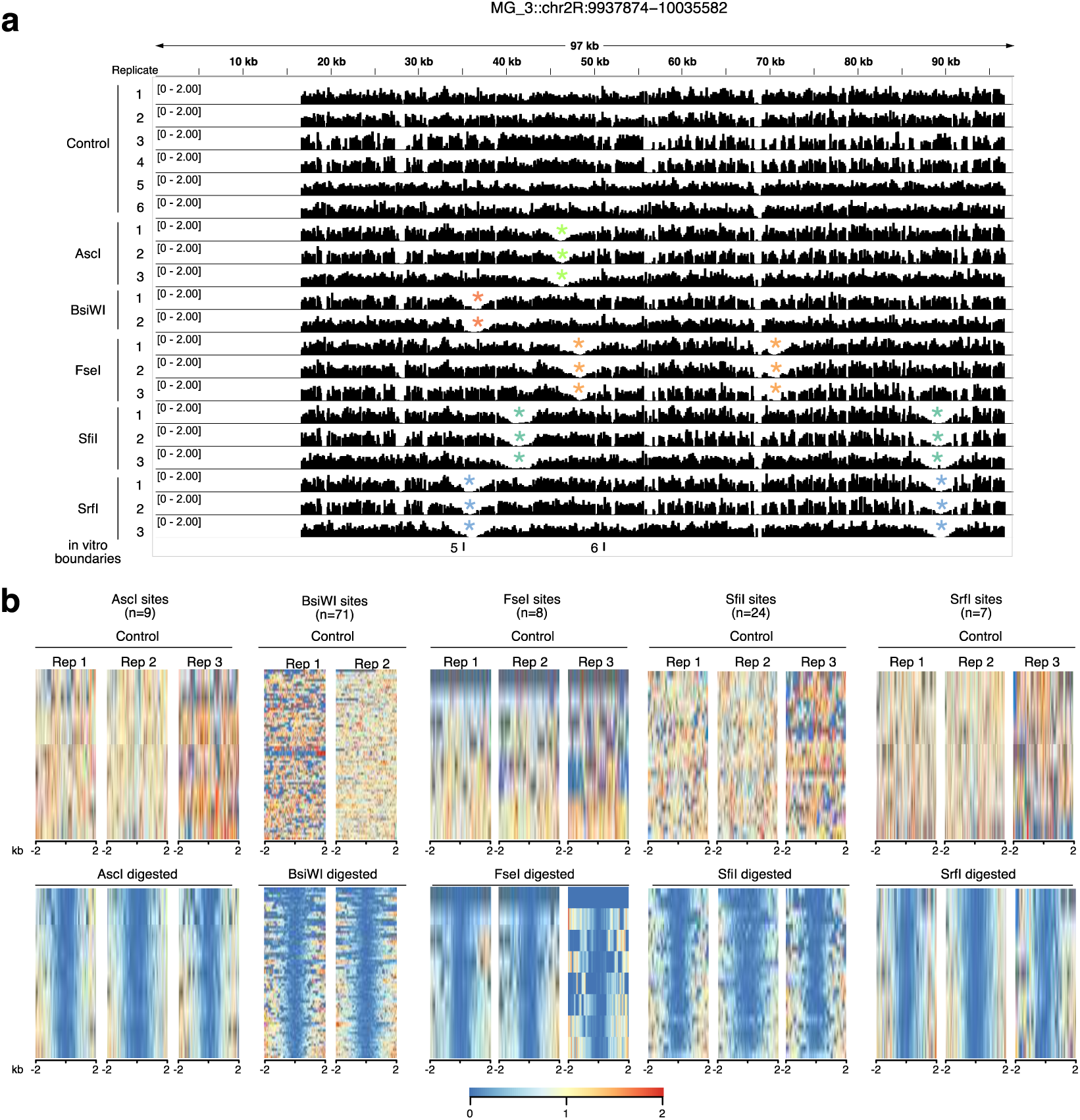
**Verification of restriction digestion efficiencies using MNase-seq data** a) MNase-seq profiles for individual replicates from restriction digestion experiments for the *eve* locus. Replicate number as well as restriction enzyme used are indicated to the left. The predicted cut site for a given restriction enzyme is indicated by colored asterisks. The boundaries are marked as a reference. b) Heatmaps representing MNase-seq coverage at 4-kb-windows centered around restriction digestion motifs for individual replicates for either control (top row) or digested (bottom row) samples. Each heatmap row represents MNase-seq coverage at an individual genomic location. The restriction enzyme, number of sites and replicate information are indicated above the heatmaps. Shared scale bar is shown at the bottom.

**Supplementary Figure S7.**
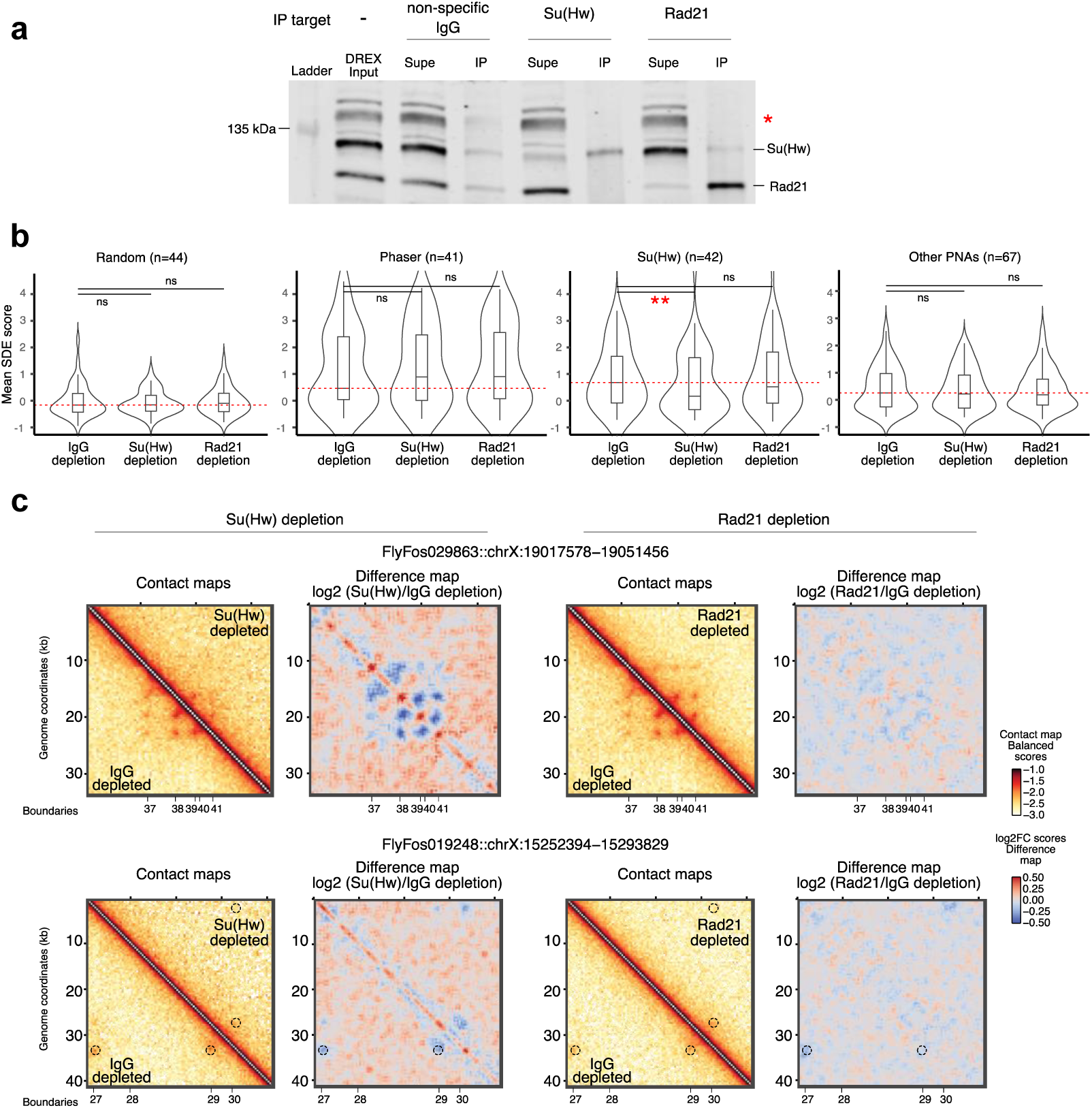
**Effects of factor depletions on nucleosome positioning and 3D genome structures** a) Western blot of replicate depletion experiments to estimate depletion efficiencies. ‘Input’ samples refer to plain DREX collected prior to depletion reactions. ‘Supe’ refers to factor-depleted DREX supernatant from the immunoprecipitation (IP) reaction while ‘IP’ refers to proteins bound to the antibody-coupled beads. Approximately 20% of immunoprecipitated material was loaded. The target of IP is indicated above the blot. The bands corresponding to the detection target are indicated on the right side of the blot with the red asterisk marking a non-specific band. The position of the135 kDa marker band is indicated to the left. b) Violin plots indicating distributions of replicate-averaged, mean periodicity SDE scores in 1-kb windows around centers of PNAs for Control, Su(Hw)-depleted and Rad21-depleted conditions grouped by PNA classes. Random positions serve as baseline reference and internal negative controls. The red dotted line indicates the median SDE-score for the corresponding control sample. Two-sided, paired Wilcoxon-signed rank tests were performed per PNA class to compare each treatment condition against the control and the resulting p-values corrected for multiple comparisons using the Benjamini-Hochberg method. The significance values are indicated as ‘ns’ (non-significant), ‘*’ (p-adjusted < 0.05) or ‘**’ (p-adjusted < 0.01). Adjusted p-values for all comparisons are provided in associated Source Data. c) 400-bp resolution Micro-C maps of FlyFos029863 and FlyFos019248 characterizing changes in contact patterns upon either Su(Hw)-depletion (left) or Rad21-depletion (right). For each region and depletion condition, interaction maps representing balanced contact frequencies for control (below the diagonal) or depleted (above the diagonal) samples are to the left, while difference maps representing log2-ratio of Depletion versus Control contact frequencies are to the right. The dotted circles represent loops referred to in the text. Called boundaries are denoted below. Scales for each heatmap type are shown to the right.

## References

1. Dekker, J., Oksuz, B.A., Zhang, Y., Wang, Y., Minsk, M.K., Kuang, S., Yang, L., Gibcus, J.H., Krietenstein, N., Rando, O.J., et al. (2026). An integrated view of the structure and function of the human 4D nucleome. Nature 649, 759–776. 10.1038/s41586-025-09890-3.

2. Misteli, T. (2020). The Self-Organizing Genome: Principles of Genome Architecture and Function. Cell 183, 28–45. 10.1016/j.cell.2020.09.014.

3. Baldi, S., Korber, P., and Becker, P.B. (2020). Beads on a string-nucleosome array arrangements and folding of the chromatin fiber. Nat Struct Mol Biol 27, 109–118. 10.1038/s41594-019-0368-x.

4. Fukai, Y.T., Kujirai, T., Wakamori, M., Kanamura, S., Yamauchi, L., Zeraati, S., Morita, S., Tanegashima, C., Kadota, M., Shirouzu, M., et al. (2025). Gene-scale in vitro reconstitution reveals histone acetylation directly controls chromatin architecture. Sci Adv 11, eadx9282. 10.1126/sciadv.adx9282.

5. Hsieh, T.H., Weiner, A., Lajoie, B., Dekker, J., Friedman, N., and Rando, O.J. (2015). Mapping Nucleosome Resolution Chromosome Folding in Yeast by Micro-C. Cell 162, 108–119. 10.1016/j.cell.2015.05.048.

6. Ricci, M.A., Manzo, C., Garcia-Parajo, M.F., Lakadamyali, M., and Cosma, M.P. (2015). Chromatin fibers are formed by heterogeneous groups of nucleosomes in vivo. Cell 160, 1145–1158. 10.1016/j.cell.2015.01.054.

7. Ohno, M., Ando, T., Priest, D.G., Kumar, V., Yoshida, Y., and Taniguchi, Y. (2019). Sub-nucleosomal Genome Structure Reveals Distinct Nucleosome Folding Motifs. Cell 176, 520–534 e525. 10.1016/j.cell.2018.12.014.

8. Li, X., and Levine, M. (2024). What are tethering elements? Curr Opin Genet Dev 84, 102151. 10.1016/j.gde.2023.102151.

9. Batut, P.J., Bing, X.Y., Sisco, Z., Raimundo, J., Levo, M., and Levine, M.S. (2022). Genome organization controls transcriptional dynamics during development. Science 375, 566–570. 10.1126/science.abi7178.

10. Willemin, A., Szabo, D., and Pombo, A. (2024). Epigenetic regulatory layers in the 3D nucleus. Mol Cell 84, 415–428. 10.1016/j.molcel.2023.12.032.

11. Bing, X., Ke, W., Fujioka, M., Kurbidaeva, A., Levitt, S., Levine, M., Schedl, P., and Jaynes, J.B. (2024). Chromosome structure in Drosophila is determined by boundary pairing not loop extrusion. Elife 13. 10.7554/eLife.94070.

12. Uhlmann, F. (2025). A unified model for cohesin function in sisterchromatid cohesion and chromatin loop formation. Mol Cell 85, 1058–1071. 10.1016/j.molcel.2025.02.005.

13. Dekker, J., and Mirny, L.A. (2024). The chromosome folding problem and how cells solve it. Cell 187, 6424–6450. 10.1016/j.cell.2024.10.026.

14. Davidson, I.F., and Peters, J.M. (2021). Genome folding through loop extrusion by SMC complexes. Nat Rev Mol Cell Biol 22, 445–464. 10.1038/s41580-021-00349-7.

15. Mirny, L.A., Imakaev, M., and Abdennur, N. (2019). Two major mechanisms of chromosome organization. Curr Opin Cell Biol 58, 142–152. 10.1016/j.ceb.2019.05.001.

16. Sultana, H., Kunar, R., and Matera, A.G. (2025). Chromatin insulators in gene regulation and 3D genome organization. Biochem Soc Trans 53, 1387–1399. 10.1042/BST20253036.

17. da Costa-Nunes, J.A., and Noordermeer, D. (2023). TADs: Dynamic structures to create stable regulatory functions. Curr Opin Struct Biol 81, 102622. 10.1016/j.sbi.2023.102622.

18. Krietenstein, N., Abraham, S., Venev, S.V., Abdennur, N., Gibcus, J., Hsieh, T.S., Parsi, K.M., Yang, L., Maehr, R., Mirny, L.A., et al. (2020). Ultrastructural Details of Mammalian Chromosome Architecture. Mol Cell 78, 554–565 e557. 10.1016/j.molcel.2020.03.003.

19. Hsieh, T.S., Cattoglio, C., Slobodyanyuk, E., Hansen, A.S., Rando, O.J., Tjian, R., and Darzacq, X. (2020). Resolving the 3D Landscape of Transcription-Linked Mammalian Chromatin Folding. Mol Cell 78, 539–553 e538. 10.1016/j.molcel.2020.03.002.

20. Galouzis, C.C., and Furlong, E.E.M. (2022). Regulating specificity in enhancer-promoter communication. Curr Opin Cell Biol 75, 102065. 10.1016/j.ceb.2022.01.010.

21. Papantonis, A., and Oudelaar, A.M. (2025). Mechanisms of Enhancer-Mediated Gene Activation in the Context of the 3D Genome. Annu Rev Genomics Hum Genet 26, 163–188. 10.1146/annurev-genom-120423-012301.

22. Kenter, A.L., Priyadarshi, S., and Drake, E.B. (2023). Locus architecture and RAG scanning determine antibody diversity. Trends Immunol 44, 119–128. 10.1016/j.it.2022.12.005.

23. Ollikainen, N., Ma, F., Braikia, F.Z., and Sen, R. (2026). Chromatin folding principles underlying the generation of antibody diversity. Mol Cell. 10.1016/j.molcel.2025.12.023.

24. Sexton, T., Yaffe, E., Kenigsberg, E., Bantignies, F., Leblanc, B., Hoichman, M., Parrinello, H., Tanay, A., and Cavalli, G. (2012). Three-dimensional folding and functional organization principles of the Drosophila genome. Cell 148, 458–472. 10.1016/j.cell.2012.01.010.

25. Dolsten, G.A., Cofer, E.M., Bing, X.Y., Brack, B., Curlin, M., Theesfeld, C.L., Troyanskaya, O.G., Levine, M.S., and Pritykin, Y. (2025). 3D chromatin structures precede genome activation in Drosophila embryogenesis. Cell Genom 5, 101002. 10.1016/j.xgen.2025.101002.

26. Maziak, N.Z., Y; Groll, F; Brown, HE; Madich, A; Kaur, Y; Harrison, MM; Zhou, J; Vaquerizas, JM 3D Genome Reorganization Foreshadows Zygotic Genome Activation in Drosophila. Nature Genetics (in press).

27. Zunjarrao, S., and Gambetta, M.C. (2025). Principles of long-range gene regulation. Curr Opin Genet Dev 91, 102323. 10.1016/j.gde.2025.102323.

28. Ing-Simmons, E., Rigau, M., and Vaquerizas, J.M. (2022). Emerging mechanisms and dynamics of three-dimensional genome organisation at zygotic genome activation. Curr Opin Cell Biol 74, 37–46. 10.1016/j.ceb.2021.12.004.

29. Ramirez, F., Bhardwaj, V., Arrigoni, L., Lam, K.C., Gruning, B.A., Villaveces, J., Habermann, B., Akhtar, A., and Manke, T. (2018). High-resolution TADs reveal DNA sequences underlying genome organization in flies. Nat Commun 9, 189. 10.1038/s41467-017-02525-w.

30. Stadler, M.R., Haines, J.E., and Eisen, M.B. (2017). Convergence of topological domain boundaries, insulators, and polytene interbands revealed by high-resolution mapping of chromatin contacts in the early Drosophila melanogaster embryo. Elife 6. 10.7554/eLife.29550.

31. Matthews, N.E., and White, R. (2019). Chromatin Architecture in the Fly: Living without CTCF/Cohesin Loop Extrusion?: Alternating Chromatin States Provide a Basis for Domain Architecture in Drosophila. Bioessays 41, e1900048. 10.1002/bies.201900048.

32. Ramasamy, S., Aljahani, A., Karpinska, M.A., Cao, T.B.N., Velychko, T., Cruz, J.N., Lidschreiber, M., and Oudelaar, A.M. (2023). The Mediator complex regulates enhancer-promoter interactions. Nat Struct Mol Biol 30, 991–1000. 10.1038/s41594-023-01027-2.

34. van Steensel, B., and Furlong, E.E.M. (2019). The role of transcription in shaping the spatial organization of the genome. Nat Rev Mol Cell Biol 20, 327–337. 10.1038/s41580-019-0114-6.

35. van Steensel, B., and Belmont, A.S. (2017). Lamina-Associated Domains: Links with Chromosome Architecture, Heterochromatin, and Gene Repression. Cell 169, 780–791. 10.1016/j.cell.2017.04.022.

35. Farrell, J.A., and O’Farrell, P.H. (2014). From egg to gastrula: how the cell cycle is remodeled during the Drosophila mid-blastula transition. Annu Rev Genet 48, 269–294. 10.1146/annurev-genet-111212-133531.

36. Hamm, D.C., and Harrison, M.M. (2018). Regulatory principles governing the maternal-to-zygotic transition: insights from Drosophila melanogaster. Open Biol 8, 180183. 10.1098/rsob.180183.

37. Seller, C.A., Cho, C.Y., and O’Farrell, P.H. (2019). Rapid embryonic cell cycles defer the establishment of heterochromatin by Eggless/SetDB1 in Drosophila. Genes Dev 33, 403–417. 10.1101/gad.321646.118.

38. Ner, S.S., and Travers, A.A. (1994). HMG-D, the Drosophila melanogaster homologue of HMG 1 protein, is associated with early embryonic chromatin in the absence of histone H1. EMBO J 13, 1817–1822. 10.1002/j.1460-2075.1994.tb06450.x.

39. Hug, C.B., Grimaldi, A.G., Kruse, K., and Vaquerizas, J.M. (2017). Chromatin Architecture Emerges during Zygotic Genome Activation Independent of Transcription. Cell 169, 216–228 e219. 10.1016/j.cell.2017.03.024.

40. Oberbeckmann, E., Quililan, K., Cramer, P., and Oudelaar, A.M. (2024). In vitro reconstitution of chromatin domains shows a role for nucleosome positioning in 3D genome organization. Nat Genet 56, 483–492. 10.1038/s41588-023-01649-8.

41. Becker, P.B., and Wu, C. (1992). Cell-free system for assembly of transcriptionally repressed chromatin from Drosophila embryos. Mol Cell Biol 12, 2241–2249. 10.1128/mcb.12.5.2241-2249.1992.

42. Langst, G., and Becker, P.B. (2001). Nucleosome mobilization and positioning by ISWI-containing chromatin-remodeling factors. J Cell Sci 114, 2561–2568. 10.1242/jcs.114.14.2561.

43. Becker, P.B. (2024). Cell-free genomics: transcription factor interactions in reconstituted naive embryonic chromatin. Biochem Soc Trans 52, 423–429. 10.1042/BST20230878.

44. Scharf, A.N., Meier, K., Seitz, V., Kremmer, E., Brehm, A., and Imhof, A. (2009). Monomethylation of lysine 20 on histone H4 facilitates chromatin maturation. Mol Cell Biol 29, 57–67. 10.1128/MCB.00989-08.

45. Volker-Albert, M.C., Pusch, M.C., Fedisch, A., Schilcher, P., Schmidt, A., and Imhof, A. (2016). A Quantitative Proteomic Analysis of In Vitro Assembled Chromatin. Mol Cell Proteomics 15, 945–959. 10.1074/mcp.M115.053553.

46. Eggers, N., and Becker, P.B. (2021). Cell-free genomics reveal intrinsic, cooperative and competitive determinants of chromatin interactions. Nucleic Acids Res 49, 7602–7617. 10.1093/nar/gkab558.

47. Baldi, S., Jain, D.S., Harpprecht, L., Zabel, A., Scheibe, M., Butter, F., Straub, T., and Becker, P.B. (2018). Genome-wide Rules of Nucleosome Phasing in Drosophila. Mol Cell 72, 661–672 e664. 10.1016/j.molcel.2018.09.032.

48. Fujioka, M., Sun, G., and Jaynes, J.B. (2013). The Drosophila eve insulator Homie promotes eve expression and protects the adjacent gene from repression by polycomb spreading. PLoS Genet 9, e1003883. 10.1371/journal.pgen.1003883.

49. Tonelli, A., Cousin, P., Jankowski, A., Wang, B., Dorier, J., Barraud, J., Zunjarrao, S., and Gambetta, M.C. (2025). Systematic screening of enhancer-blocking insulators in Drosophila identifies their DNA sequence determinants. Dev Cell 60, 630–645 e639. 10.1016/j.devcel.2024.10.017.

50. Acemel, R.D., and Lupianez, D.G. (2023). Evolution of 3D chromatin organization at different scales. Curr Opin Genet Dev 78, 102019. 10.1016/j.gde.2022.102019.

51. Ke, W., Fujioka, M., Schedl, P., and Jaynes, J.B. (2024). Stem-loop and circle-loop TADs generated by directional pairing of boundary elements have distinct physical and regulatory properties. Elife 13. 10.7554/eLife.94114.

52. Iurlaro, M., Stadler, M.B., Masoni, F., Jagani, Z., Galli, G.G., and Schubeler, D. (2021). Mammalian SWI/SNF continuously restores local accessibility to chromatin. Nat Genet 53, 279–287. 10.1038/s41588-020-00768-w.

53. Vian, L., Pekowska, A., Rao, S.S.P., Kieffer-Kwon, K.R., Jung, S., Baranello, L., Huang, S.C., El Khattabi, L., Dose, M., Pruett, N., et al. (2018). The Energetics and Physiological Impact of Cohesin Extrusion. Cell 173, 1165–1178 e1120. 10.1016/j.cell.2018.03.072.

54. Maeshima, K., Matsuda, T., Shindo, Y., Imamura, H., Tamura, S., Imai, R., Kawakami, S., Nagashima, R., Soga, T., Noji, H., et al. (2018). A Transient Rise in Free Mg(2+) Ions Released from ATP-Mg Hydrolysis Contributes to Mitotic Chromosome Condensation. Curr Biol 28, 444–451 e446. 10.1016/j.cub.2017.12.035.

55. Harpprecht, L., Baldi, S., Schauer, T., Schmidt, A., Bange, T., Robles, M.S., Kremmer, E., Imhof, A., and Becker, P.B. (2019). A Drosophila cell-free system that senses DNA breaks and triggers phosphorylation signalling. Nucleic Acids Res 47, 7444–7459. 10.1093/nar/gkz473.

56. Rowley, M.J., Nichols, M.H., Lyu, X., Ando-Kuri, M., Rivera, I.S.M., Hermetz, K., Wang, P., Ruan, Y., and Corces, V.G. (2017). Evolutionarily Conserved Principles Predict 3D Chromatin Organization. Mol Cell 67, 837–852 e837. 10.1016/j.molcel.2017.07.022.

57. Rowley, M.J., Lyu, X., Rana, V., Ando-Kuri, M., Karns, R., Bosco, G., and Corces, V.G. (2019). Condensin II Counteracts Cohesin and RNA Polymerase II in the Establishment of 3D Chromatin Organization. Cell Rep 26, 2890–2903 e2893. 10.1016/j.celrep.2019.01.116.

58. Bag, I., Chen, S., Rosin, L.F., Chen, Y., Liu, C.Y., Yu, G.Y., and Lei, E.P. (2021). M1BP cooperates with CP190 to activate transcription at TAD borders and promote chromatin insulator activity. Nat Commun 12, 4170. 10.1038/s41467-021-24407-y.

59. Mauksch, C., Zhu, Y., Velychko, T., Sagropoulou, S., Aljahani, A., Ramasamy, S., Zumer, K., and Oudelaar, A.M. (2026). FACT depletion demonstrates a role for nucleosome organization in TAD formation. Mol Syst Biol 22, 119–138. 10.1038/s44320-025-00165-7.

60. Delamarre, A., Bailey, B., Yavid, J., Koche, R., Mohibullah, N., and Whitehouse, I. (2025). Chromatin architecture mapping by multiplex proximity tagging. Mol Cell 85, 2796–2811 e2795. 10.1016/j.molcel.2025.06.018.

61. Li, H., Dalgleish, J.L.T., Lister, G., Maristany, M.J., Huertas, J., Dopico-Fernandez, A.M., Hamley, J.C., Denny, N., Bloye, G., Zhang, W., et al. (2025). Mapping chromatin structure at base-pair resolution unveils a unified model of cis-regulatory element interactions. Cell 188, 7175–7193 e7119. 10.1016/j.cell.2025.10.013.

62. Gabriele, M., Brandao, H.B., Grosse-Holz, S., Jha, A., Dailey, G.M., Cattoglio, C., Hsieh, T.S., Mirny, L., Zechner, C., and Hansen, A.S. (2022). Dynamics of CTCF- and cohesin-mediated chromatin looping revealed by live-cell imaging. Science 376, 496–501. 10.1126/science.abn6583.

63. Sabate, T., Lelandais, B., Robert, M.C., Szalay, M., Tinevez, J.Y., Bertrand, E., and Zimmer, C. (2025). Uniform dynamics of cohesin-mediated loop extrusion in living human cells. Nat Genet 57, 3152–3164. 10.1038/s41588-025-02406-9.

64. Piwko, P., Vitsaki, I., Livadaras, I., and Delidakis, C. (2019). The Role of Insulators in Transgene Transvection in Drosophila. Genetics 212, 489–508. 10.1534/genetics.119.302165.

65. Viets, K., Sauria, M.E.G., Chernoff, C., Rodriguez Viales, R., Echterling, M., Anderson, C., Tran, S., Dove, A., Goyal, R., Voortman, L., et al. (2019). Characterization of Button Loci that Promote Homologous Chromosome Pairing and Cell-Type-Specific Interchromosomal Gene Regulation. Dev Cell 51, 341–356 e347. 10.1016/j.devcel.2019.09.007.

66. Varisco, M.C., G.R; Viales, R.R; Schaub, C; Girardot, C; Forneris, M; Comeault, A; Furlong, E.E.M (2025). Heterotypic directional motifs contribute to TAD boundary function in Drosophila. bioRxiv. 10.1101/2025.09.06.674533.

67. Bhattacharya, M., Lyda, S.F., and Lei, E.P. (2024). Chromatin insulator mechanisms ensure accurate gene expression by controlling overall 3D genome organization. Curr Opin Genet Dev 87, 102208. 10.1016/j.gde.2024.102208.

68. Chathoth, K.T., Mikheeva, L.A., Crevel, G., Wolfe, J.C., Hunter, I., Beckett-Doyle, S., Cotterill, S., Dai, H., Harrison, A., and Zabet, N.R. (2022). The role of insulators and transcription in 3D chromatin organization of flies. Genome Res 32, 682–698. 10.1101/gr.275809.121.

69. Kahn, T.G., Savitsky, M., Kuong, C., Jacquier, C., Cavalli, G., Chang, J.M., and Schwartz, Y.B. (2023). Topological screen identifies hundreds of Cp190- and CTCF-dependent Drosophila chromatin insulator elements. Sci Adv 9, eade0090. 10.1126/sciadv.ade0090.

70. Li, L., Lyu, X., Hou, C., Takenaka, N., Nguyen, H.Q., Ong, C.T., Cubenas-Potts, C., Hu, M., Lei, E.P., Bosco, G., et al. (2015). Widespread rearrangement of 3D chromatin organization underlies polycomb-mediated stress-induced silencing. Mol Cell 58, 216–231. 10.1016/j.molcel.2015.02.023.

71. Ejsmont, R.K., Sarov, M., Winkler, S., Lipinski, K.A., and Tomancak, P. (2009). A toolkit for high-throughput, cross-species gene engineering in Drosophila. Nat Methods 6, 435–437. 10.1038/nmeth.1334.

72. Serizay, J., Matthey-Doret, C., Bignaud, A., Baudry, L., and Koszul, R. (2024). Orchestrating chromosome conformation capture analysis with Bioconductor. Nat Commun 15, 1072. 10.1038/s41467-024-44761-x.

73. Stansfield, J.C., Cresswell, K.G., and Dozmorov, M.G. (2019). multiHiCcompare: joint normalization and comparative analysis of complex Hi-C experiments. Bioinformatics 35, 2916–2923. 10.1093/bioinformatics/btz048.

74. Langmead, B., and Salzberg, S.L. (2012). Fast gapped-read alignment with Bowtie 2. Nat Methods 9, 357–359. 10.1038/nmeth.1923.

75. Lindenbaum, P. (2015). JVarkit: java-based utilities for Bioinformatics. 10.6084/m9.figshare.1425030.

76. Jayakrishnan, M., Havlova, M., Veverka, V., Regnard, C., and Becker, P.B. (2025). Genomic context-dependent histone H3K36 methylation by three Drosophila methyltransferases and implications for dedicated chromatin readers. Nucleic Acids Res 53. 10.1093/nar/gkaf202.

77. Heinz, S., Benner, C., Spann, N., Bertolino, E., Lin, Y.C., Laslo, P., Cheng, J.X., Murre, C., Singh, H., and Glass, C.K. (2010). Simple combinations of lineage-determining transcription factors prime cis-regulatory elements required for macrophage and B cell identities. Mol Cell 38, 576–589. 10.1016/j.molcel.2010.05.004.

78. Robinson, J.T., Thorvaldsdottir, H., Winckler, W., Guttman, M., Lander, E.S., Getz, G., and Mesirov, J.P. (2011). Integrative genomics viewer. Nat Biotechnol 29, 24–26. 10.1038/nbt.1754.

79. Casper, J., Speir, M.L., Raney, B.J., Perez, G., Nassar, L.R., Lee, C.M., Hinrichs, A.S., Gonzalez, J.N., Fischer, C., Diekhans, M., et al. (2025). The UCSC Genome Browser database: 2026 update. Nucleic Acids Res. 10.1093/nar/gkaf1250.

80. Gu, Z., Eils, R., and Schlesner, M. (2016). Complex heatmaps reveal patterns and correlations in multidimensional genomic data. Bioinformatics 32, 2847–2849. 10.1093/bioinformatics/btw313.

81. Duan, J., Rieder, L., Colonnetta, M.M., Huang, A., McKenney, M., Watters, S., Deshpande, G., Jordan, W., Fawzi, N., and Larschan, E. (2021). CLAMP and Zelda function together to promote Drosophila zygotic genome activation. Elife 10. 10.7554/eLife.69937.

82. Blythe, S.A., and Wieschaus, E.F. (2015). Zygotic genome activation triggers the DNA replication checkpoint at the midblastula transition. Cell 160, 1169–1181. 10.1016/j.cell.2015.01.050.

83. Zolotarev, N., Fedotova, A., Kyrchanova, O., Bonchuk, A., Penin, A.A., Lando, A.S., Eliseeva, I.A., Kulakovskiy, I.V., Maksimenko, O., and Georgiev, P. (2016). Architectural proteins Pita, Zw5,and ZIPIC contain homodimerization domain and support specific long-range interactions in Drosophila. Nucleic Acids Res 44, 7228–7241. 10.1093/nar/gkw371.

84. Blythe, S.A., and Wieschaus, E.F. (2016). Establishment and maintenance of heritable chromatin structure during early Drosophila embryogenesis. Elife 5. 10.7554/eLife.20148.

85. Chen, S., Rosin, L.F., Pegoraro, G., Moshkovich, N., Murphy, P.J., Yu, G., and Lei, E.P. (2022). NURF301 contributes to gypsy chromatin insulator-mediated nuclear organization. Nucleic Acids Res 50, 7906–7924. 10.1093/nar/gkac600.

86. Consortium., m., Roy, S., Ernst, J., Kharchenko, P.V., Kheradpour, P., Negre, N., Eaton, M.L., Landolin, J.M., Bristow, C.A., Ma, L., et al. (2010). Identification of functional elements and regulatory circuits by Drosophila modENCODE. Science 330, 1787–1797. 10.1126/science.1198374.

87. Wang, Q., Sun, Q., Czajkowsky, D.M., and Shao, Z. (2018). Sub-kb Hi-C in D. melanogaster reveals conserved characteristics of TADs between insect and mammalian cells. Nat Commun 9, 188. 10.1038/s41467-017-02526-9.

